# Host-associated genetic differentiation in the face of ongoing gene flow: ecological speciation in a pathogenic parasite of freshwater fish

**DOI:** 10.1101/2024.01.27.577373

**Authors:** Masoud Nazarizadeh, Milena Nováková, Jakub Vlček, Jan Štefka

## Abstract

Adaptation to varying environments, leading to population divergence, is one of the key processes of natural selection. However, its effectiveness amidst ongoing gene flow remains controversial. Our study explores this phenomenon by focusing on a tapeworm parasite (*Ligula intestinalis*), which is capable of parasitising a wide spectrum of fish species, overcoming their immunological defence and having a highly pathogenic impact. We analysed the population genetic structure, the degree of gene flow, and the level of genomic divergence between sympatrically occurring parasites from different cyprinid fish hosts. Utilising genome-wide Single Nucleotide Polymorphisms (SNPs) and transcriptome data, we investigated whether individual host species impose selection pressures on the parasite populations. Genetic clustering analyses indicated a divergence between the parasites infecting breams and those in roaches, bleaks and rudds. Historical demography modelling suggested that the most plausible scenario for this divergence is isolation with continuous gene flow. Selection analysis identified 896 SNPs under selection, exhibiting higher nucleotide diversity and genetic divergence compared to neutral loci. Transcriptome profiling corroborated these results, revealing distinct gene expression profiles for the two parasite populations. An in-depth examination of the selected SNPs and differentially expressed genes revealed specific genes and their physiological functions, as candidates for the molecular mechanisms of immune evasion and, thus, for driving ecological speciation in the parasite. This study showcases the interplay between host specificity, population demography and disruptive selection in ecological speciation. By dissecting the genomic factors at play, we gain a better understanding of the mechanisms facilitating population divergence in the presence of gene flow.

## 1 Introduction

The study of ecological speciation facilitates a comprehensive understanding of the complex interplay between evolutionary mechanisms and ecological interactions (Feder et al., 2012; Nosil et al., 2012; Rundle and Nosil, 2005). This form of speciation occurs when divergent selection pressures, associated with environmental factors, induce reproductive isolation (Nosil, 2012). In allopatric situations, reproductive isolation may arise due to accumulation of genetic variation, independently of selection pressures and adaptation to local conditions (Howard and Berlocher, 1998; Orr and Smith, 1998; Schluter, 1996). Conversely, in sympatric scenarios where populations occupy the same geographic area, reproductive isolation typically stems from reinforcement processes dampening gene flow between emerging population lineages (Bush, 1994).

Despite growing research in ecological speciation, our understanding of this process in parasites with complex life cycles remains limited (Brunner and Eizaguirre, 2016; Henrich and Kalbe, 2016; Poulin, 2011). Complex life-cycle parasites present unique opportunities for study, given their multi-host life cycles that expose them to varied ecological pressures (Combes, 2001; Poulin, 2011). Complexity arises when parasites must adapt to different hosts, each offering a distinct set of ecological challenges such as varied immune responses, habitats, and behaviours (Auld and Tinsley, 2015). Interestingly, the unique ecological conditions of parasites, such as their strict habitat selection and potential for intra-host speciation, can make sympatric speciation a more common mode of speciation for parasites compared to free-living animals (Le Gac and Giraud, 2004; McCoy, 2003). Recent studies revealed that speciation by disruptive selection for host choice plays a crucial role in sympatric speciation in parasites (de Meeûs et al., 1995; Duffy et al., 2008; Thaenkham et al., 2022). Over time, this selective pressure can lead to the formation of specialised subpopulations of parasites, each adapted to a different host. Eventually, these subpopulations may become reproductively isolated, leading to sympatric speciation (McCoy, 2003). Additionally, a hard selective sweep occurs in response to various selective pressures, such as host immune responses, drug treatments, or competition among parasites for host resources (Ebert and Fields, 2020; Le Pennec et al., 2023). When an advantageous mutation arises that better equips the parasite to deal with one of these pressures, that allele could rapidly sweep through the population. This rapid adaptation could lead to the formation of a specialised subpopulation of parasites, tuned to a specific host or set of conditions (Ebert and Fields, 2020). Despite the advances in genomic tools, there remains a gap in our understanding of how these mechanisms interplay in the context of complex life cycles and the precise genomic underpinnings that drive parasite diversification.

*Ligula intestinalis* (Cestoda) is a widespread pseudophyllidean tapeworm that serves as a valuable model organism for exploring the intricate dynamics of ecological speciation (Bouzid et al., 2008; Hoole et al., 2010; Nazarizadeh et al., 2023, 2022; Štefka et al., 2009). It possesses a complex life cycle that begins with eggs hatching in freshwater environments into coracidia larvae, which are then ingested by a copepod, the initial intermediate host. Here they develop into procercoid larvae. After ingestion of infected copepods by planktivorous fish,the second intermediate host, the larva bores through the gut into the body cavity and develops into a plerocercoid stage. The cycle is finished in a piscivorous bird, the final host, where the tapeworm reaches maturity in the bird’s gut (Dubinina, 1980). The plerocercoid phase, involving the second intermediate host, is of particular interest because this is where the parasite accumulates body mass for its subsequent maturation into an adult. By infecting numerous species of freshwater fish, plerocercoids encounter a range of ecological variables and host-specific selective pressures, such as different immune responses and hormonal manipulation of the host, which is castrated by the parasite (Dubinina, 1980; Halimi et al., 2013; Williams and Hoole, 1995). Previous genetic studies found different strains or species of the parasite, sometimes co-occurring in the same water bodies (Bouzid et al., 2008; Nazarizadeh et al., 2023; Olson et al., 2002; Štefka et al., 2009). For instance, Olson et al. (2002) identified significant genetic variations in *Ligula* populations between two sympatric fish hosts in Lough Neagh, Northern Ireland. Furthermore, recent phylogeographic research has revealed that this parasite has undergone multiple modes of speciation, including both sympatric and allopatric, resulting in its diversification into at least ten distinct evolutionary lineages across various biogeographical realms (see Figure S1 and figure 2 in Nazarizadeh et al., 2023). This ecological complexity and diverse host range provide an ideal setting for studying how speciation occurs in response to ecological factors, particularly given the parasite’s ability to adapt to different host species.

Of all ten known evolutionary lineages, Lineage A is particularly noteworthy due to its wide distribution in western Palaearctic and wide fish host spectrum, including cyprinids such as freshwater bream (*Abramis brama*), white bream (*Blicca bjoerkna*), roach (*Rutilus rutilus*), rudd (*Scardinius erythrophthalmus*), bleak (Alburnus alburnus), minnow *(Phoxinus phoxinus*), chub (*Squalius cephalus*), and crucian carp (*Carassius carassius*) (Bouzid et al., 2008; Štefka et al., 2009; Nazarizadeh et al., 2023). Interestingly, it has been documented that host preferences vary considerably in different water bodies. For example, in south-western France Loot et al. (2001) found that *L. intestinalis* primarily targets roach populations, even when other potential hosts are found in the same area. These ecological data might be indicative of emerging host-specific races of the parasite; however, a subsequent microsatellite study using 15 polymorphic loci revealed only low genetic structuring within Lineage A, with almost no variation attributable to different fish hosts across the European continent (Štefka et al., 2009). In contrast, a recent study focusing on sympatric populations from four different fish species found nonrandom distribution of mitochondrial haplotypes between the hosts, suggesting possible evolution of host specificity (Nazarizadeh et al., 2022). Considering its unique ecological character, Lineage A presents an ideal model to explore the interactions between divergent selection pressures and ecological isolation mechanisms.

Here, we aim to elucidate the mechanisms underlying possible ecological speciation in *L. intestinalis*, particularly focusing on Lineage A populations in areas where all hosts coexist in sympatry. We intend to examine the potential divergent selection pressures and ecological isolation mechanisms that might influence parasite adaptations and diversity in sympatric settings. Utilising a comprehensive approach that involves the analysis of genome-wide SNPs and transcriptome data, we strive to: (a) ascertain whether host specificity in Lineage A can potentially influence the population structure of parasites in the absence of geographical separation, and (b) identify potential genomic signatures indicative of host specialisation within identified *Ligula* populations through selection analyses. Moreover, we aspire to expand our understanding of the complex dynamics of parasite evolution, especially in the context of ecological speciation under varied host pressures. Finally, we aim (c) to study RNA transcription patterns to identify differentially expressed genes (DEGs) related to host specialisation in parasite populations by establishing a comprehensive reference transcriptome sequence.

## 2 Material and Methods

### 2.1 Sample collection

We collected 84 plerocercoid samples from five prevalent host species in Czechia: *R. rutilus*, *A. alburnus*, *B. bjoerkna*, *S. erythrophthalmus*, and *A. brama* (Nazarizadeh et al., 2022). These specimens were sourced from 10 different freshwater ecosystems throughout Czechia (see Table S1 and Figure 1) using gillnets, in the frame of hydrobiological research (see Nazarizadeh et al., 2022 for sampling details). The samples were fixed in 96% ethanol and subsequently stored at a refrigerated temperature prior to DNA extraction. For the purpose of transcriptome sequencing, 14 *Ligula* samples were taken from three fish hosts (*R. rutilus*, *B. bjoerkna*, and *A. brama)*. The tapeworms were extracted within a sterile environment, with cleanliness ensured by the use of Ultra-Pure DNAse/RNase-free water (Ambion, Austin, Texas, USA). The central segment of each tapeworm was sectioned into several 5x5 mm squares, which were then submerged in RNAlater (Ambion, Austin, Texas, USA) and maintained at 4 °C overnight to stabilise and protect the RNA, and then stored at −80 °C until RNA extraction.

**Figure 1.**
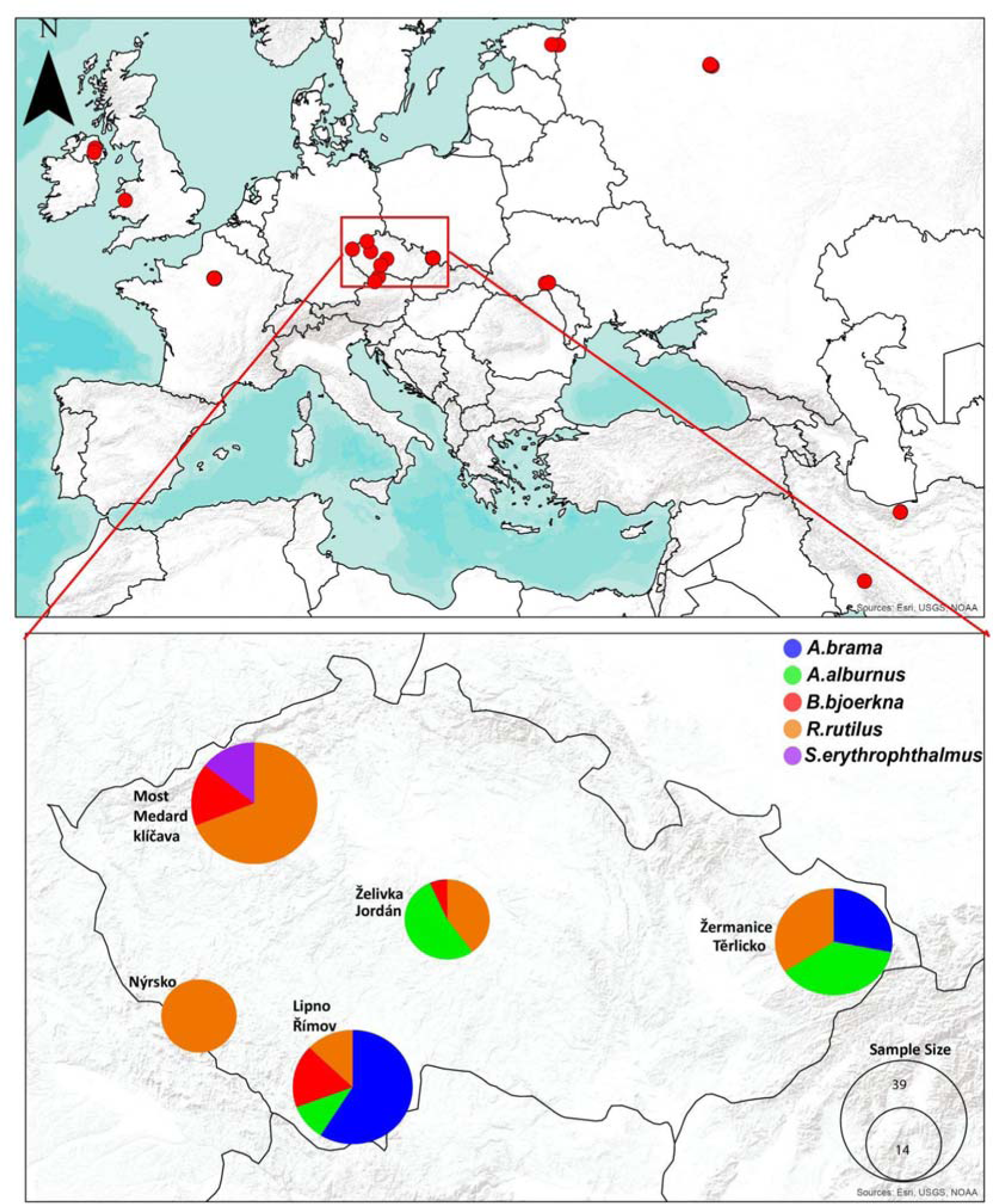
The distribution of Lineage A in *L. intestinalis* across the Palearctic (from Nazarizadeh et al., 2023, also analysed here). Inset: Sampling localities in Czechia, colour-coded to reflect the five most prevalent host species. Sampling sites in proximity to each were amalgamated into a single circle. The size of each circle correlates with the sample size.

### 2.2 DNA extraction and ddRAD library preparation

Total genomic DNA was extracted from specimens preserved in 96% ethanol using the DNeasy Blood and Tissue Kit (Qiagen). The quality and quantity of the DNA yields were verified using a 0.8% agarose gel and a Qubit 2.0 Fluorometer, respectively. Double digest restriction-site associated DNA (ddRAD) libraries were generated in accordance with the modified ddRAD protocol established by Peterson et al., (2012). We utilised the same restriction enzymes (nspI + mluCI) that were previously employed to generate ddRAD data for *L. intestinalis* (Nazarizadeh et al., 2023). All 84 samples were pooled into two libraries, which were then sequenced in two lanes to maintain uniform coverage. This process generated paired-end reads of 150 bp on an Illumina NovaSeq (Novogene UK), yielding approximately 6.7 million paired-end reads per sample. The procedures for assembling multiplexed ddRAD-seq libraries for each barcoded individual sample (Table S1), as well as the detailed purification processes, were adopted from Nazarizadeh et al. (2023).

### 2.3 Transcriptome sequencing

To compare the transcriptome profiles among different parasite populations, we selected 14 plerocercoid samples from three species of fish hosts (*R. rutilus*, *A. brama*, and *B. bjoerkna*) for transcriptome analysis. RNA extraction followed the method described by Chomczynski and Sacchi (1987), utilising the acid guanidinium thiocyanate-phenol-chloroform procedure with reagents from Invitrogen (Carlsbad, CA, USA). The RNA yield was estimated using the Qubit RNA Broad Range Assay Kit (Thermo Fisher), and its integrity was assessed with an Agilent Bioanalyzer 2100 (Agilent Technologies, USA). Subsequently, the RNA samples underwent commercial processing to be converted into cDNA libraries, followed by sequencing using the Illumina Novaseq 150 PE read technology facilitated by Novogene (UK).

### 2.4 ddRAD data assembly and SNP calling

We applied the process_radtags program from Stacks v.2.5.6 to demultiplex and filter out raw reads with low-quality or unidentified bases (Rochette et al., 2019). This phase entailed the removal of cut sites, barcodes, and adaptors from the ddRAD results. Next, we utilised FastQC (Andrews, 2010) and MultiQC (Ewels et al., 2016) to evaluate the initial quality of our raw sequencing data. To enhance our dataset, we integrated 59 ddRAD datasets from Lineage A, which were derived from an earlier population genomic study on *L. intestinalis* (Nazarizadeh et al., 2023). Following this, all paired-end reads were aligned to the *Ligula* reference genome (Nazarizadeh et al., 2024; BioProject PRJNA1055111) using the default settings of Bowtie v2.5.0 (Langmead and Salzberg, 2012). We then sorted and converted all alignments to the BAM format using Samtools v1.18 (Danecek et al., 2021), and carried out reference assembly using the ref_map.pl wrapper in Stacks for genotype calling and locus assembly. We established criteria to selectively include loci present across all populations (-p option) and in at least 80% of individuals within each population (-r), with a minor allele frequency of 3 (--min-mac) and a maximum observed heterozygosity of 80%. For further analyses focusing on population genetic structure and FST-based evaluations, we discarded SNPs in linkage disequilibrium (LD) using the --write-random-snp function in the population program. During post-processing, we used vcftools v0.1.16 (Danecek et al., 2011) to eliminate variants with extremely low (<5x) or high (800x) coverage depth, and loci with over 20% missingness. Consequently, we generated four distinct SNP matrices in VCF format as follows: **Dataset A** incorporates all samples from Lineage A (145 samples), featuring 331,589 SNPs and an average locus coverage of 21x; **Dataset B** contains 51,373 SNPs/loci LD filtering across all Lineage A individuals, boasting an average locus coverage of 18x; **Dataset C** was developed exclusively for samples from Czechia, where host species co-exist in sympatry at majority of the sampled locations (118 individuals), and it includes 333,497 SNPs, averaging a 21x locus depth of coverage; **Dataset D** comprises only unlinked SNPs from the Czech samples, including 72,696 SNPs/loci and an average locus coverage of 17x.

### 2.5 Genome-wide diversity

We used the R package snpR (Hemstrom and Jones, 2023) to compute several metrics of genetic diversity - including nucleotide diversity (Pi), standardised individual heterozygosity (Hs), expected heterozygosity (He), observed heterozygosity (Ho), the inbreeding coefficient (Fis), and the proportion of polymorphic loci (P) - across different parasite populations, utilising Dataset A which contains 331,589 SNPs. Additionally, we used the adegenet R packages (Jombart, 2008) to calculate the fixation index (Fst; Weir and Cockerham, 1984) and the genetic Nei Distance, drawing upon Dataset B which includes 51,373 SNPs.

### 2.6 Population Genetic Structure

To understand how host specificity affects population genetic structure of parasite populations, we conducted genetic clustering analyses on two datasets: all samples in Lineage A (Dataset B) and only samples from Czechia (Dataset D), aiming to compare the genetic structure of parasite populations on both large and local scales. Firstly, we performed a discriminant analysis of principal components (DAPC; Jombart et al. 2010) utilising the adegenet package in R v4.0.5. Using the K-means method, we determined the optimal number of genetic clusters based on the lowest Bayesian Information Criterion (BIC) value. Subsequently, individuals were assigned to clusters using DAPC and the optim.a.score method to establish the number of principal components to retain. We then visualised the individual memberships in each cluster through plots and showcased the clusters using Principal Component Analysis (PCA). Secondly, we employed the evolutionary clustering method in ADMIXTURE (Alexander and Lange, 2011), leveraging the parallel processing capabilities of AdmixPiPe (Mussmann et al., 2020). In this step, we conducted 20 replicates for each K value ranging from 1 to 7. The best K values were determined by the lowest cross-validation error (CV) across replicates, as suggested by Alexander & Lange (2011). The clustering was displayed using the CLUMPAK server (http://clumpak.tau.ac.il/). Lastly, we used fineRADstructure and RADpainter v.0.2 (Malinsky et al., 2018) to explore the population genetic structure based on the nearest-neighbour haplotype. We ran the Stacks2fineRAD.py script from the fineRADstructure package to calculate the distribution of alleles and SNPs per locus, and the amount of missing data per individual, allowing a maximum of 10 SNPs per locus and limiting individual missingness to 25% during the conversion of the haplotype file to the RADpainter format. Considering the sensitivity of ddRAD to batch effects caused by minor differences between libraries at the size selection step, we examined the potential impact of missing data on any library-based structure. We utilised the fineRADstructure pipeline with default settings but increased the burn-in iterations to 200,000, with 1,000,000 iterations sampled at 1000 intervals. We evaluated convergence by assigning individuals to populations across multiple independent runs, reviewing the plots for the MCMC output of parameter values to ensure consistent convergence on Bayesian posterior distributions, and obtaining effective parameter sample sizes (above 100) by lengthening the duration of each chain. To plot the co-ancestry heatmap, we utilised the “FinestructureLibrary.R” function in the fineRADstructure package (Malinsky et al., 2018).

### 2.7 Coalescent analysis of Speciation modelling

To better understand the impact of isolation and migrations on the population dynamics of Lineage A, we utilised a coalescent-based approach examining gene flow and demographics. Four speciation models were proposed, and their corresponding demographic models were applied to two parasite populations, as defined by the analyses of population genetic structure. The first speciation scenario was defined as allopatric speciation, characterised by complete geographic isolation without gene flow. The second speciation model accounted for post-divergence gene flow through isolation after migration (primary contact), while the third speciation model tested post-divergence with secondary contact, which led to recent gene flow between the two parasite populations. Finally, the fourth scenario hypothesised isolation with continuous gene flow between parasite populations (Figure 2).

**Figure 2.**
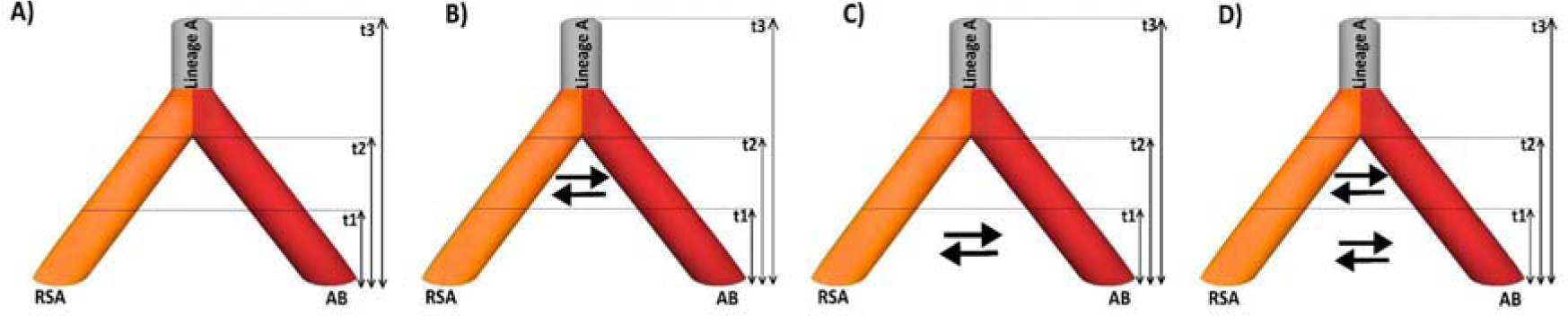
The four speciation scenarios testing the gene flow and demography patterns in Lineage A. A) Represents the allopatric speciation model with complete geographic isolation indicated by no gene flow between parasite populations from *R. rutilus*, *S. erythrophthalmus*, and *A. alburnus* (RSA) and parasite populations from *A.brama* and *B. bjoerkna (*AB). B) Depicts the model with post-divergence gene flow through primary contact. C) Shows post-divergence speciation with secondary contact leading to recent gene flow. D) Illustrates the scenario with continuous gene flow, as indicated by multiple gene flow arrows between populations over time.

We calculated the composite likelihood of our observed data within a particular model using the site frequency spectrum (SFS) and the simulation method provided by fastsimcoal2 (Excoffier & Foll, 2011; Excoffier et al., 2013). For the simulations, we selected 25 samples for each genetic group representing the highest probability of assignment to their specific genetic cluster (i.e., we excluded highly admixed parasite populations between two genetic clusters; see the Results section). In the genotype calling process for our 50 samples, we utilised the Stacks software, adopting a stringent selection criterion to substantially reduce the incidence of missing data. This approach involved retaining only those loci that were consistently present across all individuals in both populations. Then, using the easySFS Python script (Coffman et al., 2016; Gutenkunst et al., 2009), we generated a folded joint SFS, considering a single SNP for each locus to reduce the effects of linkage disequilibrium. We determined the fixed population size in our analysis from the nucleotide diversity (π) and the mutation rate (μ) per site for each generation, as indicated by the formula Ne = (π / 4μ) (Nei, 1987). Nucleotide diversity (π) for parasite populations from *R. rutilus*, *A. alburnus*, and *S. erythrophthalmus* (RSA), as well as those in *A. brama* and *B. bjoerkna* (AB), was estimated from both polymorphic and non-polymorphic loci using Stacks. The average mutation rate per site per generation (2.89 × 10^-9) and demographic history events were obtained from Nazarizadeh et al., 2023 (see figure S1). The time of change in demographic events was estimated as a model parameter and allowed to range from 1 to 300,000 years (generations). Each model was run in 100 replicates, with each replicate consisting of 1,000,000 simulations, for the calculation of the composite likelihood, 50 expectation-conditional maximisation (ECM) cycles, and a stopping criterion of 0.001 (Papadopoulou and Knowles, 2015). We used an information-theoretic model selection approach based on the Akaike’s information criterion (AIC) to determine the probability of each model given the observed data. After the maximum likelihood was estimated for each model in every replicate, we calculated the AIC scores (Thomé and Carstens, 2016). AIC values for each model were rescaled (DAIC) by calculating the difference between the AIC value of each model and the minimum AIC obtained among all competing models. Point estimates of the different demographic parameters for the best-supported model were selected from the run with the highest maximum composite likelihood. Finally, we calculated confidence intervals of parameter estimates from 100 parametric bootstrap replicates by simulating SFS from the maximum composite likelihood estimates and re-estimating parameters each time (Excoffier et al., 2013).

### 2.8 Isolation by Distance

To investigate a potential correlation between genetic distance and geographic distance among parasite populations, we considered Dataset B to perform an isolation by distance (IBD) analysis using the Mantel test in the adegenet package in R (Jombart, 2008; Mentel, 1967). This analysis required calculating the Nei genetic distances and the geographic distances (measured in kilometres between population locations), which were then correlated through the mantel.randtest function. To reduce the impact of host specificity on genetic distance, we extended the analysis to various genetic structures, reinforced by a Monte Carlo simulation with 999 permutations for stronger statistical inference. Following this, we sought to clarify the nature of the observed correlations, determining if they illustrated a continuous or a patchy distant cline of genetic differentiation. To do this, we employed a 2-dimensional Kernel density estimator through the kde2d function found in the MASS package (Ripley et al., 2013) in R v.3.6.2, enhancing our grasp of the spatial distribution of genetic variation across the area.

### 2.9 Gene flow among parasite populations

To evaluate the gene flow among parasite populations in sympatric hosts, we used the unlinked SNPs data set from Czechia (Dataset D) to reconstruct the interactions among these groups. First, the Treemix v.1.12 (Pickrell and Pritchard, 2012) tool was utilised to inspect the gene flow amongst populations in a phylogenetic context. Based on allelic frequency data, a Maximum Likelihood (ML) tree was constructed and past migratory interactions between populations were inferred. Individual migrations (n) were hypothesised and calculated separately. The second ddRAD dataset was selected and divided into six clusters based on host specificity. This analysis was conducted using information from five diverse host-derived parasite populations. A range of migration events (m) from 1 to 6 was evaluated (1+ the total number of populations). The best model was determined by considering the covariance related to each migration event, and the stability of the tree structure was verified through bootstrap replicas, generated from SNP blocks consisting of 1,000 units. The results obtained from Treemix were depicted utilising the popcorn package in R.

Moreover, the contemporary gene flow among parasite groups originating from various hosts was analysed using the Bayesian inference incorporated in BayesAss, facilitated by the BA3SNP software (Mussmann et al., 2019). This analysis encompassed 10 million iterations, discarding the initial 1 million steps, and data were collected at intervals of 1,000 steps. To enhance the precision of the evaluation, cross-validation was performed on the SNP datasets. Adjustments were made in the -a and -f parameters to achieve acceptance rates ranging between 20 and 60% for allele frequencies and inbreeding coefficients, respectively (Mussmann et al., 2019).

### 2.10 Genomic signatures of host specific selection

To test for evidence of natural selection associated with host specificity, we analysed Dataset C to identify loci under selection between two distinct parasite populations in sympatric hosts. These populations exhibit marked host specificity, as evidenced by their genetic structure. To accurately identify loci under directional selection, we used multiple selection analyses and adopted a strategy that relies on the convergence of results from several analytical methods (Tsumura et al., 2012). We employed four well-established tests to detect outlier loci: the PCA-based method outlined in pcadapt v4.3.3 (Privé et al., 2020), the FST frequency method detailed in Outflank v2 (Whitlock and Lotterhos, 2015), the Bayesian method that focuses on allele frequency variations as detailed in BayeScan v2.1 (Foll and Gaggiotti, 2008), and the variance analysis in haplotype frequencies conducted using the hapFLK v1.4.

The first outlier detection method, utilised via the R package pcadapt (v4.3.3; Luu et al., 2017), conducts an individual-based genome scan grounded in PCA. This method does not rely on prior assumptions about population groupings, thus avoiding the need to force admixed individuals into predefined populations. The initial pcadapt run was performed using 20 PCs (K=20). The ideal number of PCs to retain for later tests was determined following Cattell’s rule, as described by Luu et al. (2017). The second outlier method used the Outflank r package to determine the distribution of FST for neutral loci, which was then used to assign q-values to each locus to detect outliers that may be due to spatially heterogeneous selection. In this analysis, we set the ‘number_of_samples’ parameter to 5 (equal to the number of populations sampled), the ‘LeftTrimFraction’ to 0.08, the ‘RightTrimFraction’ to 0.30, and maintained the default setting for the Hmin parameter (0.1). The initial threshold for calculating q-values was 0.05, as set by default (Whitlock and Lotterhos, 2015)

Additionally, we utilised the BayeScan v2.192 to pinpoint loci undergoing divergent selection, a process grounded in the variations in allele frequencies across distinct populations. SNP loci with a false discovery rate (FDR) below 0.05 were chosen as outlier SNPs. The analysis included an initial 20 pilot runs, each with 5000 iterations, succeeded by a main phase of 500,000 iterations with a burn-in period of 250,000 steps to guarantee convergence. The default prior odds value of 10 was retained throughout. Based on their alpha values, the loci were categorized: loci significantly above zero were regarded as under directional selection, while those below zero were considered to be under balancing selection (Moore et al., 2014), with the rest being categorized as neutral. The resulting data, which included FST values, were imported into R software using the BayeScan package for further analysis following Geweke’s diagnostic method, known for its efficient convergence diagnostics and outlier detection. This yielded a comprehensive list and tally of outliers, aiding in the visualization and interpretation of selection patterns across the examined populations.

Furthermore, the HapFLK software was employed for a haplotype-based analysis on regions potentially experiencing selection, with the local haplotype cluster (K) set to 20 and the number of iterations increased to 20, balancing accuracy and computational time. Essentially, HapFLK expands on FST-based analysis, pinpointing genomic areas with notable haplotype divergence between individuals from selected populations, while acknowledging the population structure. This method is efficient in identifying recent selective sweeps differentiating the populations under study. Moreover, regions were identified as potential selection areas if at least two successive SNPs showed a nominal p-value less than or equal to 0.05. To ascertain the random appearance of regions fulfilling this criterion, a resampling test was conducted, where SNP positions were shuffled 1000 times to determine the probability of observing sequences with a specific number of consecutive SNPs meeting the defined criteria. Regions with a resampling p-value under 0.05 were retained for additional analysis. Loci identified as outliers by the four methods were deemed “potential outliers” and were presented through Venn diagrams utilizing the “VennDiagram” package in R (Chen and Boutros, 2011).

Finally, we estimated FST, Tajima’s D, per locus absolute divergence (dXY), and nucleotide diversity (pi) using the R package PopGenome (Pfeifer et al., 2014). We then plotted both FST and dXY values against pi. We expect dXY to be significantly higher for outlier loci compared to neutral loci in a scenario of divergence with gene flow, assuming a sufficient amount of time has elapsed since the initial divergence (Cruickshank and Hahn, 2014). Moreover, dXY is expected to elevate in highly differentiated regions, resulting in a positive correlation between FST and dXY. Furthermore, we employed the “KStest” function from the GSAR R package (Rahmatallah et al., 2017) to perform the Kolmogorov–Smirnov D test, comparing Tajima’s D values between outlier and neutral loci. This helped ascertain if the distributions of Tajima’s D values from putative outlier loci were statistically different from those of putative non-outlier loci. A significant difference (p < 0.05) in distributions would imply that outlier loci have been subjected to distinct evolutionary pressures (i.e., selection, genetic drift, gene flow) compared to the rest of the genome.

### 2.11 Differentially Expressed Genes (DEGs) associated with host specificity

14 RNA-seq samples were used to compare transcriptome differences between two parasite populations detected by population genetic structure. Trimmomatic v0.33 (Bolger et al., 2014) was used to discard low quality reads from all samples. Following trimming, paired-end reads (Table S2) were mapped to the *L. intestinalis* reference genome (BioProject: PRJNA1055111) using STAR (Dobin et al., 2013). To eliminate reads that mapped to multiple loci and also to sort and convert the filtered SAM files to BAM format, we used Samtools v1.18 (Danecek et al., 2021). Furthermore, the mapped reads counted for genomic features using featureCounts v1.06 in the Subread package (Liao et al., 2014). Table S3 presents mapped reads per sample replicate and gene model. To normalise the raw counts, we applied the Trimmed Mean of M values (TMM) method. The edgeR (Robinson et al., 2010) was used to estimate differential expression of genes between parasite populations based on their host specificity. Additionally, we applied the criteria of an absolute log-fold change greater than 1 or less than -1 and a p-value less than 0.05 to identify DEGs. Finally, using the plotMDS function in edgeR, we plotted the first two principal components to evaluate the general similarities and differences among all the transcriptomes.

### 2.12 Functional annotation of loci under selection and DEGs

We implemented a two-step strategy to annotate the selected loci. In the first step, we identified outliers in the *Ligula* reference genome and retrieved the relevant annotation details using the Integrative Genomics Viewer (IGV) (Robinson et al., 2011). Next, for SNPs located in non-gene regions of the *Ligula* genome, we took into account a 201 bp sequence - comprising 100 bp both upstream and downstream of the SNP from the reference genome sequences. Using OmicsBox v3 (BioBam, 2019), we performed a comprehensive functional annotation of these sequences. Initially, nucleotide sequences (201 bp) were blasted to the NCBI non-redundant nucleotide (nt) database Pruitt et al., 2007) through BLASTX. For each search, we annotated the primary hit with the highest total score, maintaining an E-score of 10-5 for BLASTX and 10-15 for BLASTN, and insisting on a search coverage of more than 70%. We further enriched our annotations using the integrated InterProScan (Jones et al., 2014) module for domain-based information and mapped our data to the EggNOG database to provide broader functional insights (Huerta-Cepas et al., 2019).

Gene Ontology (GO) annotations for *L. intestinalis* genes were sourced from Nazarizadeh et al., 2024. We conducted functional enrichment analysis using the GSEA function in the clusterprofiler R package (Wu et al., 2021), applying the Benjamini-Hochberg false discovery rate adjustment with a 0.05 threshold. We used the DOSE R package (Yu et al., 2014) to create dot plots and enrichment maps for the highlighted genes. GOs were summarized via REVIGO (revigo.irb.hr) for DEG sets, adopting a 0.4 threshold and using SimRel for similarity (Supek et al., 2011).

## 3 Results

### 3.1 Genomic diversity

We compared the genetic characteristics of various parasite populations from different fish host species within *Ligula intestinalis* Lineage A (Table 1). Populations found in *R. rutilus* hosts located in Czechia, France, Ireland, and Ukraine showed the highest nucleotide diversity (0.013) and expected heterozygosity (0.131). In contrast, parasites from *S. orientalis* hosts in Iran and *A. brama* hosts in Czechia, Estonia, and Russia displayed the lowest nucleotide diversity, with a pi value of 0.0122. Interestingly, parasites in *S. orientalis* hosts in Iran had the highest observed heterozygosity (Ho) of 0.172, notably higher than the expected heterozygosity. However, parasites in *B. bjoerkna* hosts in Czechia exhibited the lowest Ho, marked at 0.089. Additionally, the highest proportions of polymorphic loci (P), valued at 1.02, were identified in parasite populations of *R. rutilus* and *S. erythrophthalmus* hosts. Conversely, the lowest proportion, 0.89, is observed in parasite populations of *S. orientalis* hosts (Table 1).

**Table 1.** Genetic diversity metrics of parasite populations in sampled hosts and localities. The table shows nucleotide diversity (Pi), standardized individual heterozygosity (Hs), expected heterozygosity (He), observed heterozygosity (Ho), Inbreeding coefficient (Fis), and the proportion of polymorphic loci (P).

| Parasite population | Hosts | Locality | N | Pi | Hs | He | Ho | Fis | P |
| --- | --- | --- | --- | --- | --- | --- | --- | --- | --- |
| Lineage A | *R. rutilus* | Czechia,France,Ireland, Ukraine | 73 | 0.013 | 1.02 | 0.131 | 0.118 | 0.106 | 0.93 |
|  | *A. brama* | Czechia, Estonia, Russia, | 28 | 0.012 | 0.944 | 0.120 | 0.115 | 0.116 | 0.65 |
|  | *A. alburnus* | Ukraine, Czechia | 21 | 0.012 | 1.00 | 0.126 | 0.116 | 0.107 | 0.65 |
|  | *B. bjoerkna* | Czechia | 10 | 0.011 | 0.997 | 0.119 | 0.108 | 0.089 | 0.48 |
|  | *S. erythrophthalmus* | Czechia | 5 | 0.012 | 1.02 | 0.11<br>7 | 0.11<br>8 | 0.102 | 0.40 |
|  | *S. orientalis* | Iran | 4 | 0.012 | 0.89 | 0.11<br>0 | 0.10<br>3 | 0.172 | 0.35 |
|  | *C. carassius* | Ukraine | 1 | - |  | - | - | - | - |
|  | *P. Phoxinus* | United Kingdom | 1 | - |  | - | - | - | - |

Further analysis revealed the minimum Nei genetic distance as 0.006 between parasite populations of *A. brama* and *B. bjoerkna*. *R. rutilus* demonstrates the lowest genetic distance to *S. erythrophthalmus* and *A. alburnus*, marked at 0.007 and 0.008, respectively. The Fst values provide insights into the genetic differentiation among these populations, with higher values indicating increased differentiation. Notably, parasite populations in *A. brama* and *A. alburnus* demonstrate the highest differentiation, evidenced by Fst value of 0.076. On the other end, *R. rutilus* and *S. erythrophthalmus* populations revealed minimal differentiation, as seen from Fst value of 0.001, indicating negligible genetic variation between them (Table S2).

### 3.2. Population genetic structure

The DAPC analysis for Dataset B (spanning all geographic locations) and Dataset D (specific to Czechia) consistently identified two primary genetic clusters within Lineage A. This finding was reinforced by the K-means method, with the best partitioning at a K value of 2, as indicated by the lowest BIC score for these datasets. In both datasets, the primary DAPC axis, DAPC 1, accounts for 39.3% and 45.1% of the genetic variance, respectively. This axis divides the populations into two distinct clusters: the first includes parasite populations from *A. brama* and *B. bjoerkna*, while the second comprises populations from a variety of other species, including *S. erythrophthalmus*, *R. rutilus*, *S. orientalis*, *P. phoxinus*, and *C. Carassius*. The secondary axis, contributing to a smaller 8.9% of the total variance, distinguishes the parasite populations in *S. orientalis* and *A. alburnus* from those in *R. rutilus* and *S. erythrophthalmus*. The results of admixture analysis from Datasets B and D align well with the DAPC analysis, identifying an optimal K value of 2, as indicated by the cross-validation error (CV, Figure S2). On a large scale, parasite populations in Lineage A are divided into two genetic clusters, mirroring the local scale where all host species coexist (Figure 3A and Figure 3B). The first cluster contains parasite populations from *A.brama* and *B. bjoerkna*, while the second groups together populations from *R. rutilus*, *A. alburnus*, and *S. erythrophthalmus*. Furthermore, the analysis showed that parasite populations from *B. bjoerkna* are significantly admixed, with a 46%– 90% assignment to the *R. rutilus* parasite population (Figure 3C).

**Figure 3.**
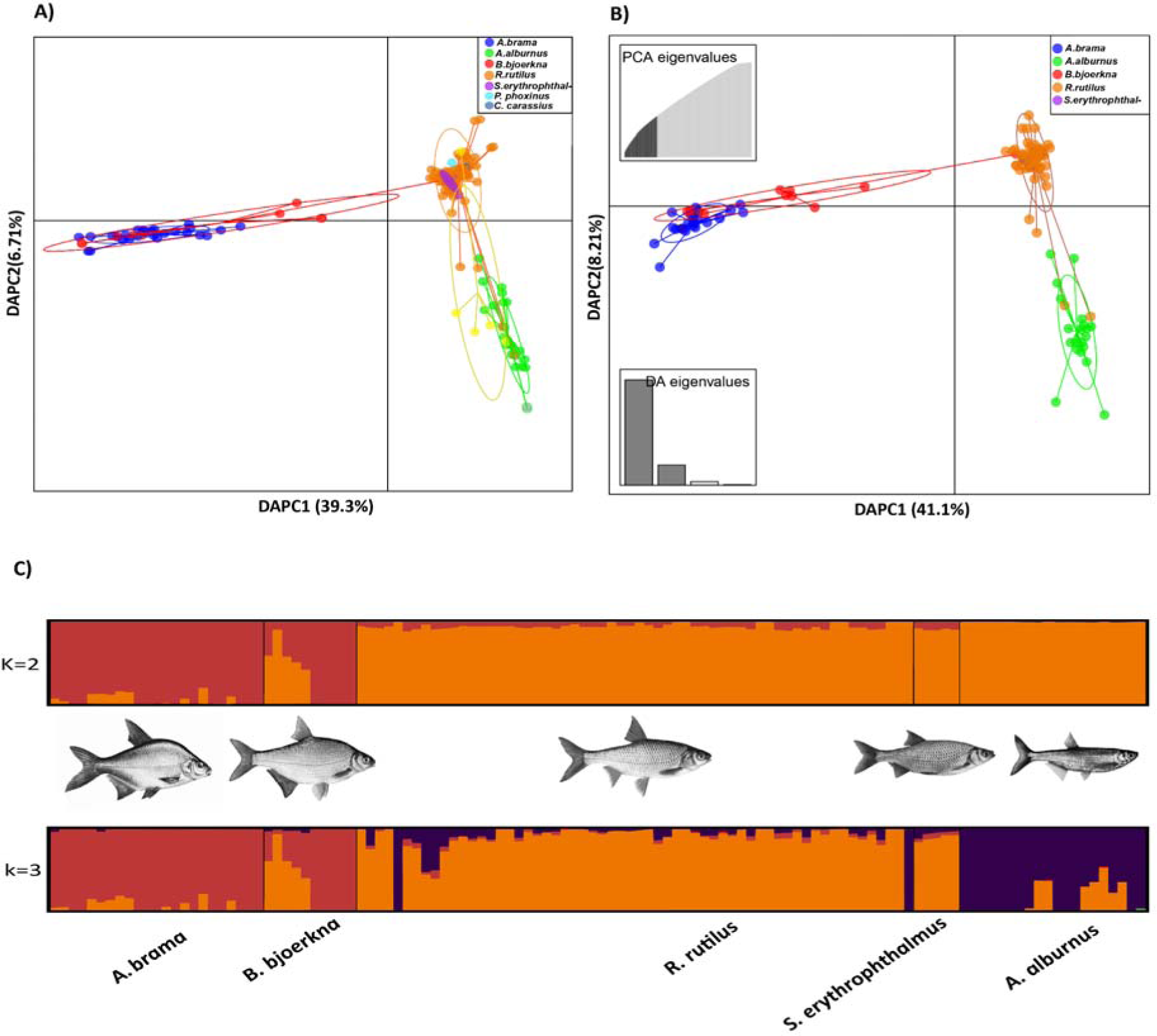
Genetic structure of parasite populations within Lineage A across different host species. A) DAPC analysis of datasets B (global, 148 samples) and D (Czechia-specific, 118 samples) reveals two primary genetic clusters, with primary axis (DAPC 1) accounting for 39.3% and 45.1% of variance, respectively. This axis predominantly segregates populations from *A. brama* and *B. bjoerkna* from other species such as *S. erythrophthalmus*, *R. rutilus*, *S. orientalis*, *P. phoxinus*, and *C. carassius*. The secondary axis highlights differentiation between *S. orientalis* and *A. alburnus* versus *R. rutilus* and *S. erythrophthalmus*, contributing to 8.9% of total variance. B) Admixture analysis is consistent with DAPC findings, particularly noting the significant admixture in *B. bjoerkna* populations, shown with a 46%–90% genetic overlap with the *R. rutilus* parasite population.

Additionally, the fineRADstructure analysis validated the clustering results with K=2, demonstrating significant support in the dendrogram. This analysis identified the parasite population in *A. alburnus* as a sub-cluster stemming from the populations in *R. rutilus* and *S. erythrophthalmus*. It also highlighted a notable level of shared co-ancestry between the parasite populations in *A. brama* and *B. bjoerkna*. However, this genetic cluster exhibits a lower level of shared co-ancestry with other parasite populations, both on a broad and local scale (Figure 4). In line with the admixture results, one sample from *B. bjoerkna* was clustered with the parasite populations in *R. rutilus*. The Mantel test results further confirmed the absence of a significant correlation between genetic distance and geographic location within Lineage A parasite populations. This suggests that geographical distances do not hinder gene flow at the intra-population level (Figure S3).

**Figure 4.**
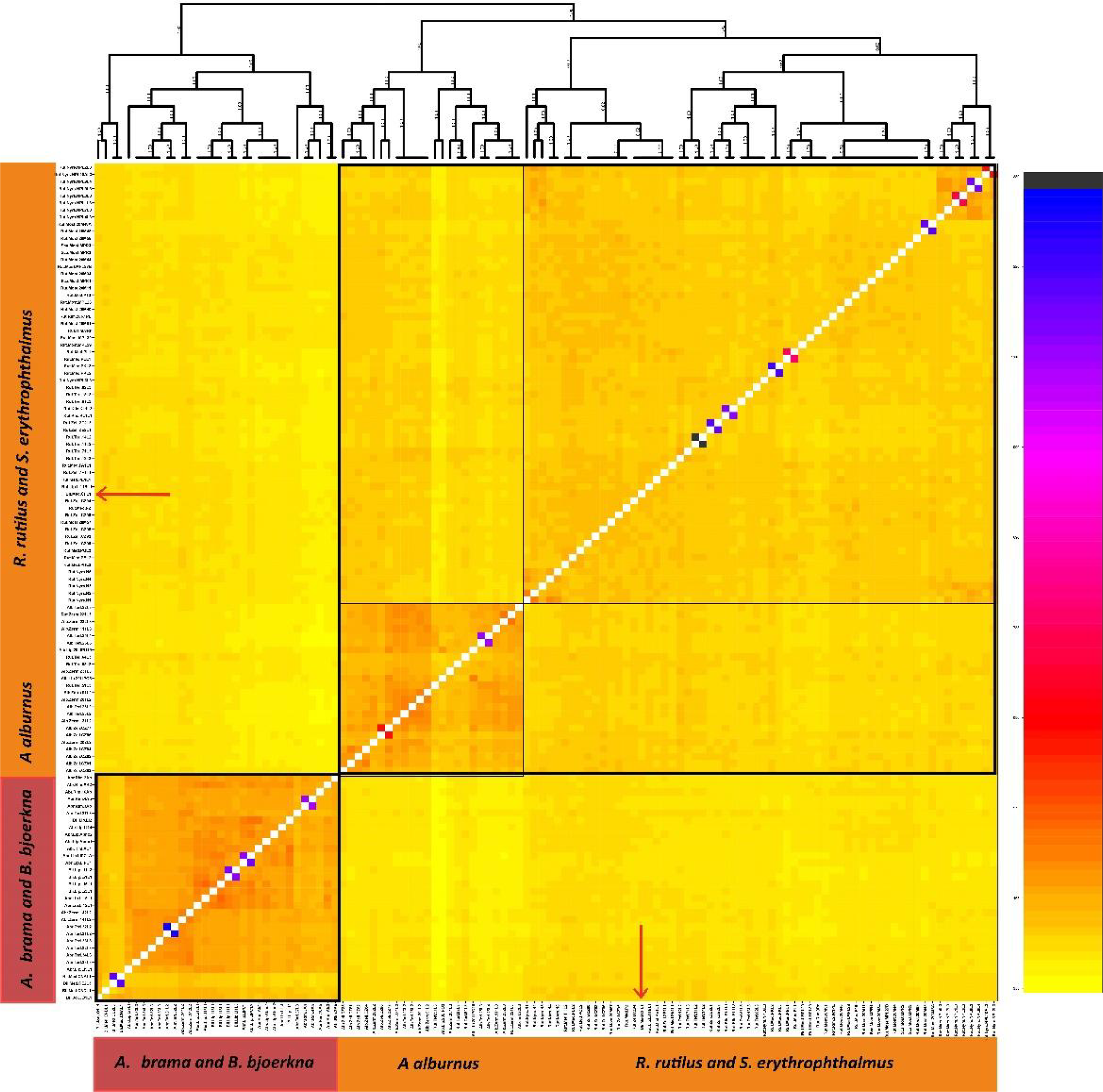
Co-ancestry plot and dendrogram derived from the fineRADstructure evaluation of *Ligula* populations in Czechia. Co-ancestry coefficients transition from low (yellow) to high (blue), indicating the extent of recent shared ancestry between a focal individual (on the vertical axis) and all other individuals included in the study (on the horizontal axis). The red arrow highlights a single sample of *Ligula* from the host *B. bjoerkna* (a host switch), which is nested in a genetic cluster from *R. rutilus* and *S. erythrophthalmus*.

### 3.2 The divergence modelling of parasite populations

Coalescent analysis for species modeling showed the most probable scenario to be Isolation with ongoing gene flow, where gene flow rates have recently risen. The change in the rate of gene flow was inferred to have occurred about 68000 generations ago. However, these time estimates should be interpreted with caution as we fixed the first split to the upper bound of 300,000 generations ago (corresponding roughly to the time since when two parasite populations probably diverged from each other, Nazarizadeh et al 2023). Splitting times may be even more recent if there was a significant time lag between gene tree and species tree. The second most likely model, with a ΔAIC of 242, suggested recent gene flow starting 925 generations ago, representing secondary contact. Models involving primary contact showed very close likelihoods (ΔAIC 2106), while the model excluding gene flow was a poor match for the data (ΔAIC 1241). These findings indicate speciation occurred amidst gene flow.However, having only two migration matrices is a strong simplification of the speciation process (Table S3).

### 3.3 Gene flow among parasite populations in different hosts

The unrooted species tree derived from the Treemix analysis identified two primary genetic groups, corroborating the findings from both the admixture and fineRADstructure analyses. Notably, the data are best explained by a model that considers two migration events. Parasite populations in *A. brama* and *B. bjoerkna* demonstrated a close genetic relationship, forming a sister lineage distinct from other parasite groups. The most prominent migration events were observed among the parasite communities in *A. alburnus*, *R. rutilus*, and *S. erythrophthalmus*. Additionally, a gene flow with a lower weight was detected between *B. bjoerkna* and *S. erythrophthalmus* populations (Figure 5A). The results of the contemporary gene flow indicated that most of the gene flow originated from populations within the same host species. While there was some gene flow between different hosts, populations in *A. brama* and *B. bjoerkna* symmetrically contributed to the pool of exogenous allelic variants. Similarly, a noticeable amount of balanced gene flow was observed from the parasite populations in *R. rutilus* to those in *A. alburnus* and *S. erythrophthalmus*. Moreover, gene flow was detected between populations in *A. brama* and *B. bjoerkna* and those in *A. alburnus*, *R. rutilus*, and *S. erythrophthalmus* (Figure 5B).

**Figure 5.**
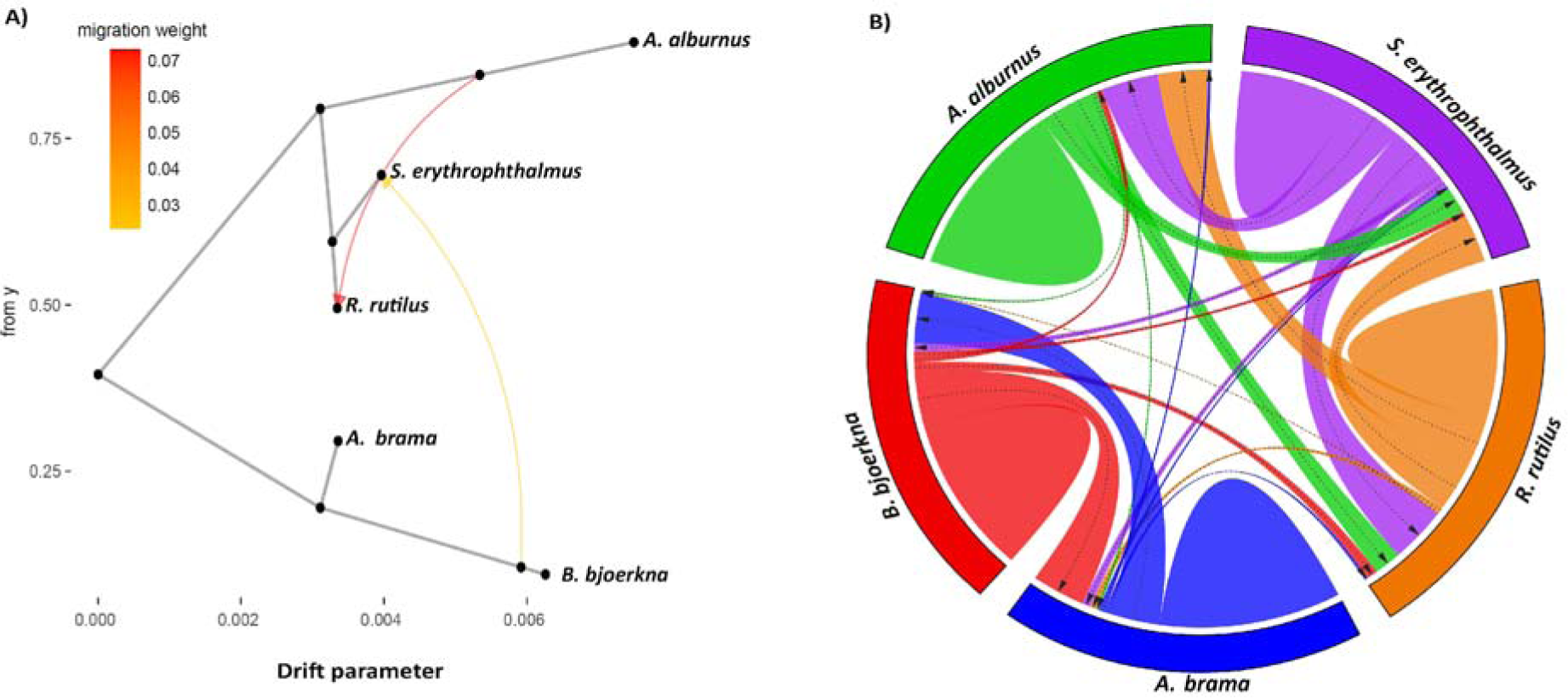
Gene flow analysis of *Ligula* populations in Czechia. (A) Species tree derived from Treemix shows two principal genetic clusters with significant gene flow among *A. alburnus*, *R. rutilus*, and *S. erythrophthalmus*. (B) analysis of BayesAss reveals contemporary gene flow among parasite populations in different hosts, particularly originating from *R. rutilus*, and underscores the contributions of *A. brama* and *B. bjoerkna* to various host populations.

### 3.4. Detection of SNPs under divergent selection

Selection analyses were conducted to identify loci potentially under divergent selection in two parasite populations structured within sympatric hosts. A total of 3,004 outlier loci, putatively under selection, were identified using PCAdapt, OutFLANK, BAYESCAN, and HapFLK. The least conservative method, BAYESCAN, identified the highest number of outliers with 2,367 SNPs, PCAdapt detected 1,743 SNPs, whilst OutFLANK and HapFLK were the most conservative, pinpointing 1,384 and 1,256 loci, respectively. Notably, all four methods collectively detected 896 SNPs (29.8% of the total SNPs), marking them as the putatively adaptive dataset (Figure S4). Most of the commonly detected loci across the methods were found in the 10 longest scaffolds of the *Ligula* genome (Figure 6). Out of 896 outlier loci identified through four methods, 156 were directly associated with the coding sections of 65 gene models in the *Ligula* genome. Considering potential gaps in the genome annotation, a second method was applied, extending the search to 100 bp before and after each outlier. This approach revealed 37 additional outliers, sharing homology with genes from closely related tapeworm species.

**Figure 6.**
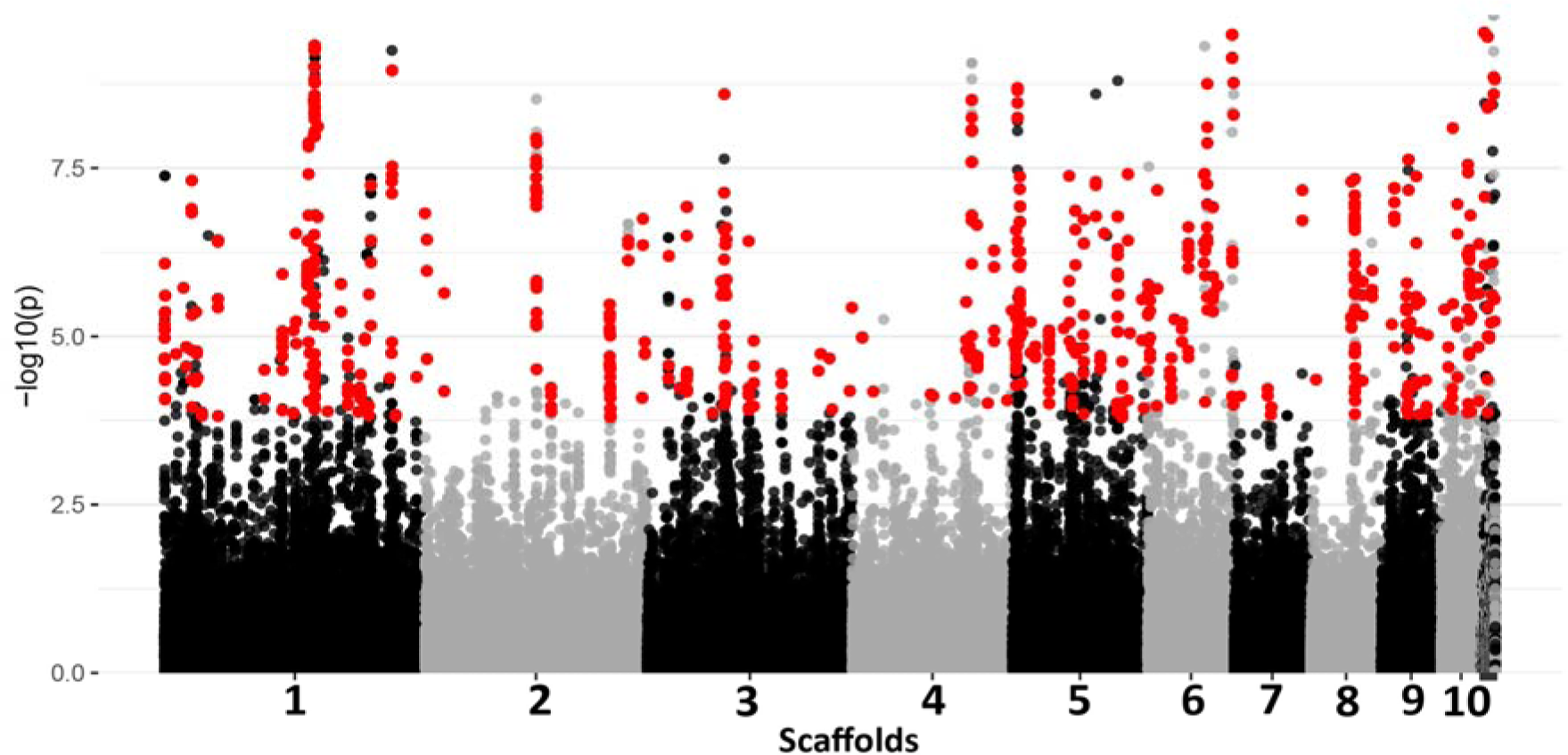
A consensus Manhattan plot displaying results from four genome-wide selection analyses: PCAdapt, Outflank, BAYESCAN, and HapFLK. This plot presents the 10 longest scaffolds of the *Ligula* genome (additional scaffolds are also included in the plot and are represented collectively due to their shorter lengths; Nazarizadeh et al., 2024). SNPs marked in red indicate those under selection between two parasite populations found in sympatric hosts. The y-axis represents -log10 (p) values.

A significant positive correlation was observed between the values of dXY and nucleotide diversity for both neutral and outlier loci. Outlier loci predominantly displayed a higher dXY compared to neutral loci and exhibited lower to moderate nucleotide diversity in contrast to neutral loci (Figure 7A). Similarly, dXY showed a positive correlation with FST, and outlier loci displayed a relatively higher FST compared to neutral loci (Figure 7B). Notably, the distributions of Tajima’s D revealed significant negative values for putative outlier in comparison to non-outlier loci (P value < 0.01, Figure S5).

**Figure 7.**
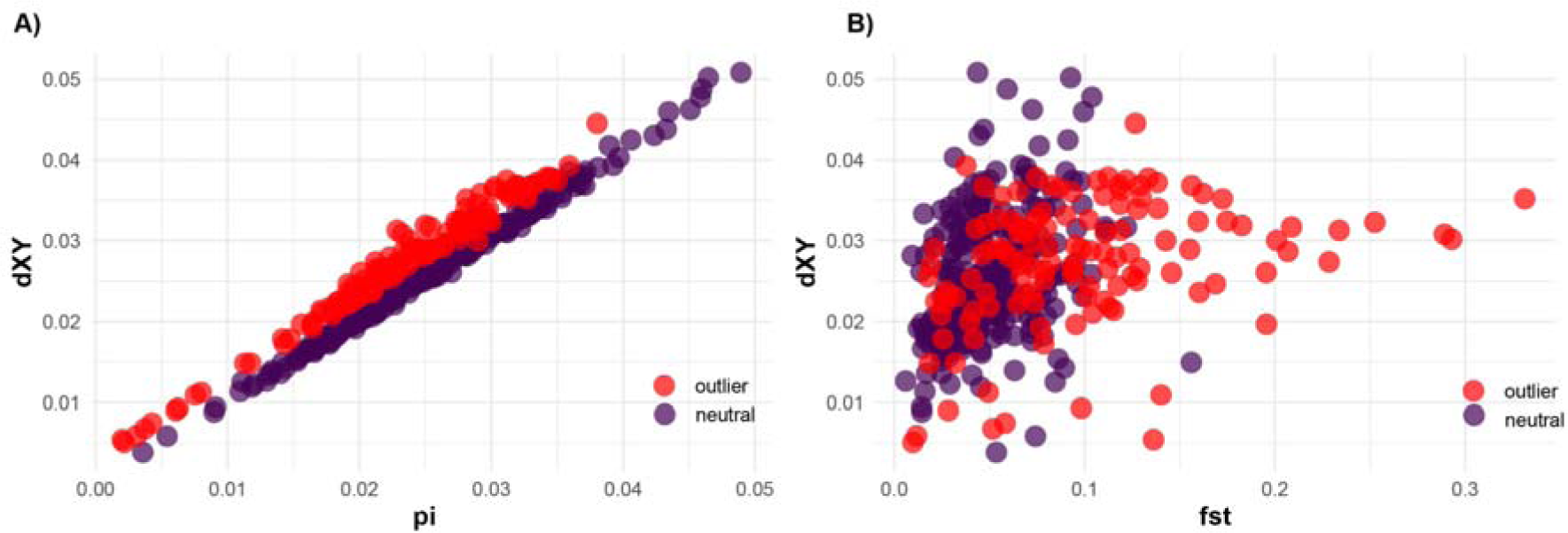
Comparative analysis of genome-wide variations between putative outlier and non-outlier loci. A) Correlation between dXY and pi distinguishing outlier from neutral loci. B) Patterns of dXY distribution in association with FST, highlighted for both outlier and neutral loci.

### 3.5 Differential gene expression

Gene expression analysis of 14 transcriptome samples derived from three different hosts in a sympatric setting, identified 993 differentially expressed genes (DEGs): 556 were up-regulated and 367 down-regulated in the parasite populations of *R. rutilus* compared to *A. brama* and *B. bjoerkna* Figure (8A and 8B). This pattern is consistent with the results of the genetic structure of the parasite populations. The RNA expression profiles showed that parasites in two of the hosts, *A. brama* and *B. bjoerkna*, had similar transcriptome patterns. In contrast, parasites in *R. rutilus* showed a distinct transcriptomic profile (Figure 8A). Furthermore, the parasite population in *R. rutilus* differed from those in *A. brama* and *B. bjoerkna* in PCA, with the first dimension accounting for 32.11% of the observed variance (Figure 8C).

**Figure 8.**
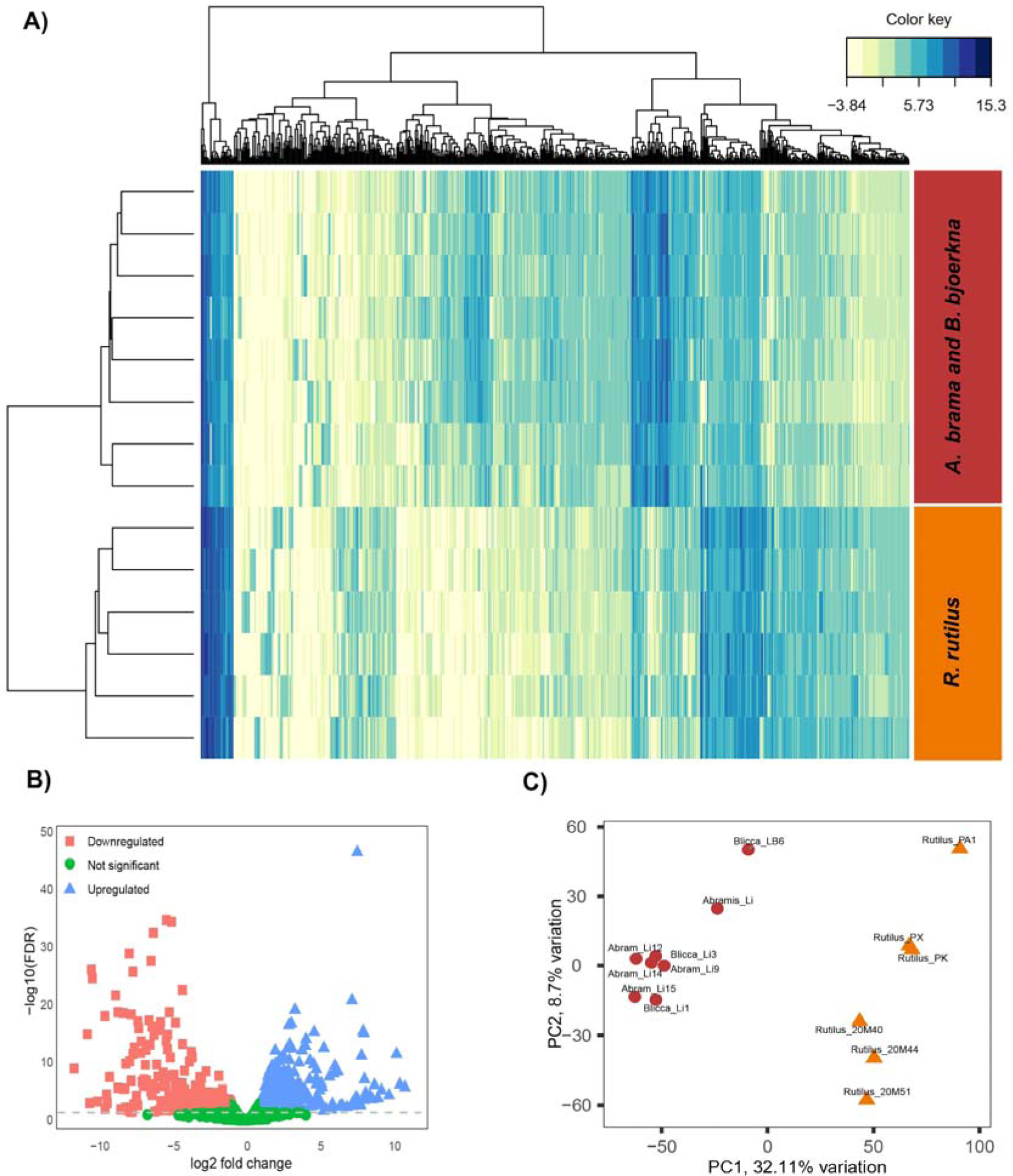
Differential gene expression in *L. intestinalis* samples obtained from three host species living in sympatry. (A) Heatmap illustrates the differential expression of genes across parasite populations in different host species. (B) Volcano plot displaying upregulated (red rectangles), downregulated (blue triangles), and non-differentially expressed transcripts (green dots) across hosts *(R. rutilus*, *A. brama,* and B. *bjoerkna*). (C) PCA showcasing the clustering of sample replicates, with relative variances detailed across PC1 and PC2.

### 3.6 Gene Ontology Patterns in DEGs and SNPs under selection

The results of the Gene Ontology analysis revealed significant variations in gene enrichment in parasite populations across different host environments. In the parasite population from *R rutilus*, the Biological Process (BP) ontology was highlighted by dominant terms such as “nervous system process” with 15 genes and “proteolysis” with 13 genes, indicating a possible focus on neural activities and protein degradation. Regarding Cellular Components (CC), genes associated with “extracellular space” and “endoplasmic reticulum” were particularly prominent, suggesting interactions between cellular organelles and the external environment. The Molecular Function ontology (MF) highlights the significant presence of “hydrolase activity” in this environment, indicating central enzymatic activities (Figure 9A). In contrast, in *L. intestinalis* populations from *A. brama* and *B. bjoerkna*, the BP ontology showed “signalling” with 15 genes, indicating important communication and regulatory pathways. The CC ontology was dominated by “nucleoplasm” with 35 genes, indicating potentially enhanced nuclear activities or regulations. “Cytoplasmic vesicles” also played an important role, indicating important intracellular transport mechanisms. Within the MF ontology, there was a notable focus on “RNA binding” and “Catalytic activity acting on DNA”, indicating essential molecular interactions and potential DNA manipulations (Figure 9B).

**Figre 9.**
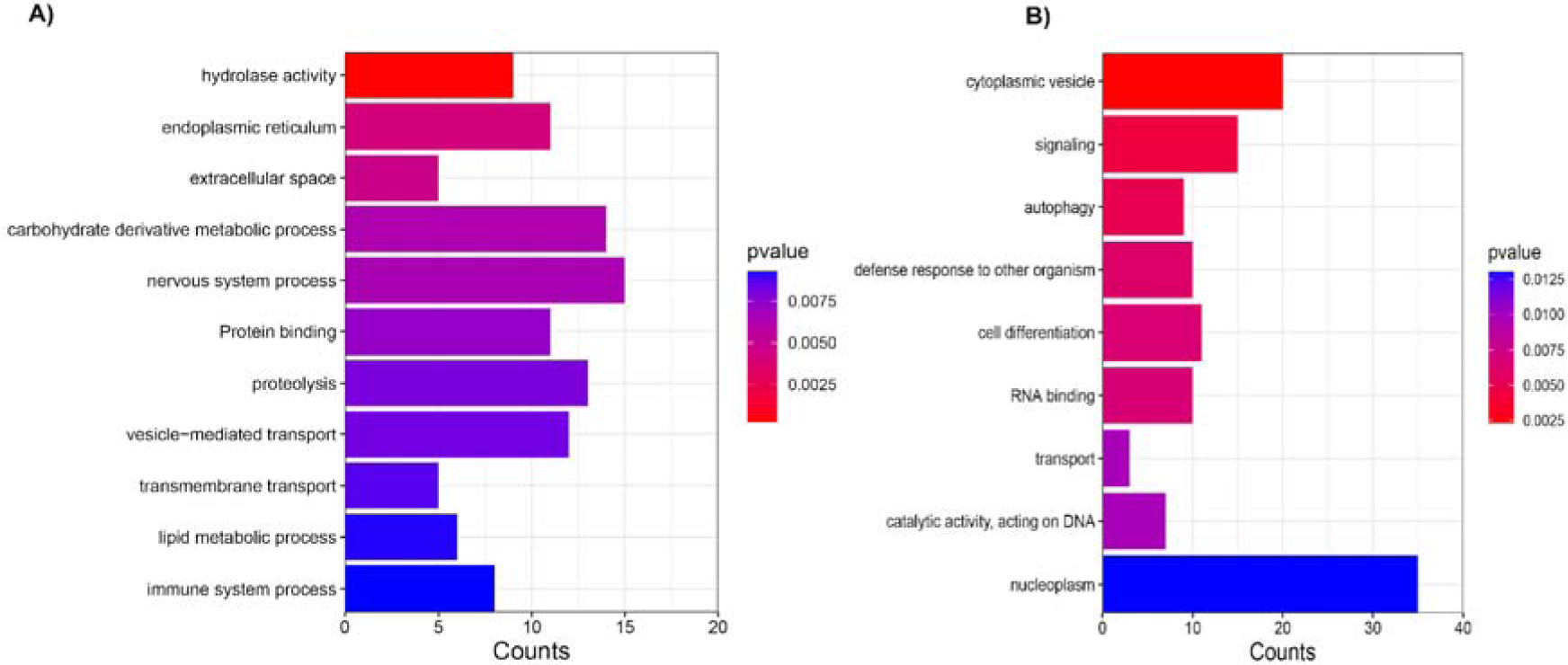
Differential Gene Ontology (GO) term enrichment in *L. intestinalis* populations parasitising different hosts. (A) Overrepresented GO terms in the parasite population from *R. rutilus*, highlighting key biological processes, cellular components, and molecular functions with the respective gene counts. (B) Overrepresented GO terms in parasite populations from *A. brama* and *B. bjoerkna* hosts, showcasing the distinct adaptive molecular responses and interactions within these environments. The y-axis represents the number of genes associated with each GO term.

The ddRAD-derived putative outlier data revealed a comprehensive profile of GO term enrichment within *L. intestinalis* populations. Most notably, the BP ontology had 140 terms that emphasised processes such as the cell junction organisation, autophagy, DNA integration, response to abiotic stimulus, generation of precursor metabolites and energy, Regulation of signalling, immune system processes and ammonium ion metabolic process (Figure S6). The enrichment for cellular components included 74 terms that emphasised cellular structures such as the cytosol, lipid droplet, sarcolemma and plasma membrane (Figure S7). The ontology for molecular functions, on the other hand, was the most diverse with 52 terms, highlighting functionalities such as cAMP binding, ether hydrolase activity, protein binding and catalytic activity (Figure S8). A comparative analysis with previously evaluated transcriptome data revealed a common enrichment of GO. These common terms from proteolysis, nucleolysis, immune system processes, signalling, autophagy, cytosol, and hydrolase activity highlight the consistent molecular and cellular activities central to *L. intestinalis* populations across different analytical platforms and datasets. Overall, five genes were identified as common between the ddRAD selection analyses and differential gene expression analysis.

## 4 Discussion

The interplay between ecological pressures and natural selection in shaping host specificity is a critical factor in the evolutionary dynamics of parasites (Simmond et al., 2020; Nikolakis et al., 2022; Bellis et al., 2022). Ecological pressures, such as environmental conditions and the availability of hosts, can greatly influence the range of organisms a parasite can infect (Cháves-González et al 2022). This creates a selective environment where certain traits are favoured, leading to adaptations in host specificity (Clark and Clegg, 2017). In the present study, we have employed genome-wide SNPs and transcriptome data to unravel the complexities of the intraspecific population structure within a single evolutionary lineage of *L. intestinalis* tapeworm. This study places a special focus on the parasite’s interactions with its second intermediate hosts, thus shedding light on how ecological factors play a significant role in driving speciation between closely related parasite populations. Our research was propelled by the intriguing observation of two distinct parasite populations coexisting in sympatry. Notably, these two clusters exhibited a high level of concordance with two different groups of host species, suggesting a subtle yet significant interplay between host specificity and the evolutionary trajectory of the parasite. Our study suggests that the genetic divergence between the two parasite clusters is indicative of speciation with ongoing gene flow, where disruptive selection facilitates reproductive isolation. Direct association between ecological divergence and reproductive isolation is a keenly sought phenomenon in specialist organisms, including parasitic plants (Giraud et al., 2010) phytophagous ladybird beetles (Matsubayashi et al., 2017), parasitic wasps (Stelinski and Liburd, 2005), leaf mites (Skoracka et al., 2013) and phytophagous insects (Berlocher and Feder, 2002; Funk et al., 2002). However, only a few studies explored this link using genomic data and in a framework including ongoing gene flow (Hume et al., 2018; Mateus et al., 2013; Villacis-Perez et al., 2021). Below, we delve into the results of our investigation, specifically examining the role of ecological speciation in the evolution and diversification of this parasite.

### 4.1 Population genetic structure and divergence with ongoing gene flow

Analysis of genetic structure using genome-wide SNP data revealed that *L. intestinalis* Lineage A is divided into two primary clusters within its Palearctic distribution. At this intraspecies level, distinct differences were observed in the parasite populations of *A. brama* and *B. bjoerkna* compared to those in *R. rutilus*, *S. erythrophthalmus*, *A. alburnus*, *P. phoxinus*, and *C. carassius*. This pattern was also evident on a more local scale within Czechia, where these host species were sampled in sympatry, highlighting the role of host specificity in the parasite’s evolution. These findings align with our previous study that revealed a non-random shared haplotype between five individuals of parasite populations in *A. brama* and *R. rutilus* (Nazarizadeh et al., 2022). Yet, earlier investigation involving more individuals showed a high number of shared mtDNA haplotypes and an indistinguishable population structure among all parasite populations in Lineage A (Bouzid et al., 2008). This result indicates that while mtDNA provides valuable insights into broad evolutionary patterns and phylogeography, it may be less effective in detecting subtle, microevolutionary changes and ecological divergence that occur over short timescales or in complex population structures. In contrast, genome-wide SNP and nuclear marker analyses offer a more comprehensive and detailed view of these processes.

Ecological divergence often interacts with gene flow, the extent of which varies with dispersal distance (Räsänen & Hendry, 2008). In our study, we observed genetic divergence among parasites from two related genetic clusters, yet we also identified notable gene flow between them. Our Treemix analysis indicated a significant migration event that suggests interbreeding between these groups. This was further supported by admixture analysis, which identified hybrid individuals among the two parasite clusters. Additionally, within one of these clusters, encompassing the fish hosts *A. alburnus*, *R. rutilus*, and *S. erythrophthalmus*, we detected another significant gene flow event. Because these specialist parasite populations have demonstrated high divergence in host use, our results clearly demonstrate that the ecological divergence has been well maintained by natural selection even in the presence of gene flow. Our results are in line with the contemporary perspective of sympatric speciation in the genomic era, in which gene flow is almost pervasive between recently diverged sister lineages (Marques et al. 2019).

We focused on distinguishing between historical secondary admixture and ongoing gene flow in the *Ligula* system using two approaches, analysis of genetic diversity in outlier loci and demographic analysis of various admixture scenarios. Analysis of 896 outlier loci p revealed positive correlation between dXY and nucleotide diversity suggesting that as populations split, genetic variation within them increases, hinting at a continued gene flow (Cruickshank and Hahn, 2014). This is further corroborated by the finding that outlier loci, which are likely under strong selective pressure, exhibit a higher degree of divergence (as indicated by higher dXY values) compared to neutral loci. Additionally, these outlier loci displayed lower to moderate nucleotide diversity, suggesting that selection is acting on these loci to a greater extent, limiting their genetic variation (Nosil et al., 2009). The correlation of dXY with FST and the significant negative values of Tajima’s D in outlier loci reinforce the notion that selection is driving specific parts of the genome to diverge, even as gene flow persists (Cruickshank and Hahn, 2014). This pattern aligns with the second approach, using coalescent species modelling, where gene flow does not preclude the emergence of distinct populations, as differential selection pressures can lead to significant divergence in key genomic regions, setting the stage for the evolution of new evolutionary species.

Recent studies in various host-parasite systems show complex patterns of gene flow and reproductive isolation in parasites (Leder et al., 2021; Momigliano et al., 2017; Simmonds et al., 2020; Teske et al., 2019). These studies highlight the potential influence of disruptive selection in the process of speciation, leading to the divergence of a single ancestral population into groups that specialise in different habitats and hosts. In the present study, the result of species modelling revealed that the most plausible evolutionary scenario for the divergence between parasite genetic clusters is speciation through isolation with migration. This involves recurrent migration after divergence, as illustrated in Figure 2a. This notion supports the increasing evidence that isolation with ongoing gene flow is a likely mechanism for speciation during ecological divergence (Nosil, 2012; Rundle and Nosil, 2005). The divergence of these two parasite clusters likely occurred in the Chibanian period (Middle Pleistocene, according to Nazarizadeh et al., 2023), possibly in tandem with the colonisation of and host shifting among various cypriniform hosts. This period, characterised by significant climatic and environmental shifts, might have led to different subgroups within a population diverging due to varying selective pressures (Walker et al., 2019). Here, ecological disruptive selection is a key driver of ecological speciation, as it enhances the fitness of each population within its respective host groups. Notably, such divergence, driven by local adaptation, can occur even among populations in close geographical proximity, leading to sympatric speciation. In such scenarios, segments within a parasite population begin to specialise in exploiting different host species within the same ecosystem. For instance, one subgroup of a parasitic species might adapt to thrive with a specific host, exploiting unique physiological or immunological traits of that host, while another subgroup might specialise in a completely different host species.

### 4.2 Candidate loci and transcriptome genes involved in adaptation to host

The study of gene expression variation within and between species has long been a subject of interest, particularly in discerning the influences of gene plasticity and natural selection (Mathieu-Bégné et al., 2022; Romero et al., 2012). If significant genetic differentiation and selection-related outlier SNPs are detected, it implies that changes in gene expression might be attributed to evolutionary adaptations, rather than just to plastic responses to varying environments (Mathieu-Bégné et al., 2022). In this study, we explored the genomic differentiation between two parasite populations living in sympatric hosts over different time scales by comparing genomic outlier loci indicative of long-term adaptation with DEGs representing short-term acclimation. Although our transcriptome data did not cover all parasite populations from the five hosts studied, our results were consistent with genome-wide SNP analyses, revealing clear separate profiles of gene expression between parasite populations from *R. rutilus* and those in *A. brama* and *B. bjoerkna*.

Furthermore, our findings include five genes showing a direct correlation between outlier loci and DEGs, either in terms of gene identity or physical proximity, along with notable functional parallels. Of these five DEGs, ANN02916 and ANN012081, are associated with reverse transcription activities. Aligning with a prior research by Nazarizadeh et al. (2024), ANN02916 was identified as significantly downregulated during the transition from the larval to the adult stage in *L. intestinalis*. The reverse transcription process involves DNA sequences in an organism’s genome derived from RNA, which is a key process in retroviruses and retrotransposons (Hughes, 2015). Additionally, our study sheds light on the biological processes associated with DNA integration through the annotation of outlier SNPs. These findings suggest that DNA integration and activities related to reverse transcription are pivotal in determining the host specificity and immune evasion strategies of the parasite. Such mechanisms are crucial not only across different host groups but also between various life stages, including larvae and adults (Nazarizadeh et al., 2024). These mechanisms contribute to genetic variability and local adaptation, which are essential for the survival and evolutionary success of the parasite within their host environments.

The tree other genes, ANN07147, ANN10664, and ANN13534, identified as significant in both SNPs under selection and DEGs play critical roles in biological functions linked to the endoplasmic reticulum (ER) membrane and transmembrane transport. Both processes may be important for the survival and host specificity of parasites, the membrane is key for carrier-mediated transport within the tapeworm tegument, greatly influencing the chemical modification of absorbed substances and offering protection against the host’s digestive enzymes. Notably, this membrane includes a glycocalyx layer made up of oligo- or polysaccharide chains attached to external lipid and protein elements. This layer is fundamental for binding various substances, such as inorganic ions and larger organic molecules including host enzymes. While some of these enzymes remain active on the worm’s surface, facilitating contact digestion, others are bound in a non-active form, possibly serving as a defence mechanism to prevent the parasite from being digested by the host. Additionally, our results highlight the biological function of glycosylation, potentially linked to the glycocalyx layer of the tegument. Studies have shown that glycosylation inhibitors like tunicamycin can substantially reduce the incorporation of galactose into the tapeworm’s tegument and carcass (Hildreth et al., 1997; Izvekova et al., 2021). This reduction highlights the importance of glycosylation in maintaining the surface glycocalyx of the tapeworm (Hildreth et al., 1997). Moreover, the presence of glucose transporter homologues in *Taenia solium* suggests a link between glucose absorption through the tegument and glycosylation processes (Rodríguez-Contreras et al., 1998). The tegument’s external limiting membrane is coated with carbohydrate-rich polyelectrolytes, indicating the presence of glycosylation and its potential role in the tapeworm’s membrane structure and function (Lumsden, 1975). This glycosylation is not only crucial for the tapeworm’s nutrient absorption but also plays a significant role in host-parasite interactions. The functional importance of glycolipids in *Spirometra erinaceieuropaei* is a prime example (Yanagisawa et al., 1999). Additionally, the presence of glycoconjugates in the tegument of *Taenia taeniaeformis* may contribute to the parasite’s resistance to host digestive processes (Olson et al., 2000).

In conclusion, using whole-genome genotyping and transcriptome analyses we demonstrated how host-specialisation leads to sympatric genetic differentiation in spite of continued gene flow. We proposed disruptive host-mediated selection on genes involved in immunological or physiological interaction with the host as the driver of ecological speciation in this pathogenic parasite of freshwater fish. This study provides an example of a frequently anticipated, but rarely observed, ecological phenomenon of speciation by host specificity.

## Acknowledgements

We would like to thank colleagues who helped with field sampling (namely Anna Mácová and Roman Hrdlička). The work was supported by a grant from the Czech Science Agency (GA19-04676S). Computational resources were provided by the project “e-Infrastruktura CZ” (e-INFRA CZ LM2018140) supported by the Ministry of Education, Youth and Sports of the Czech Republic.

## Author contributions

JŠ initiated the research topic. All authors participated in field collecting. MNov. performed laboratory analyses. MNaz. analysed the data under supervision of JŠ and JV. MNaz. and JŠ drafted the paper with input from JV and MNov.

## Notes

### Competing Interest Statement

The authors have declared no competing interest.

