## Supplemental Information for "Host-associated genetic differentiation in the face of ongoing gene flow: ecological speciation in a pathogenic parasite of freshwater fish"

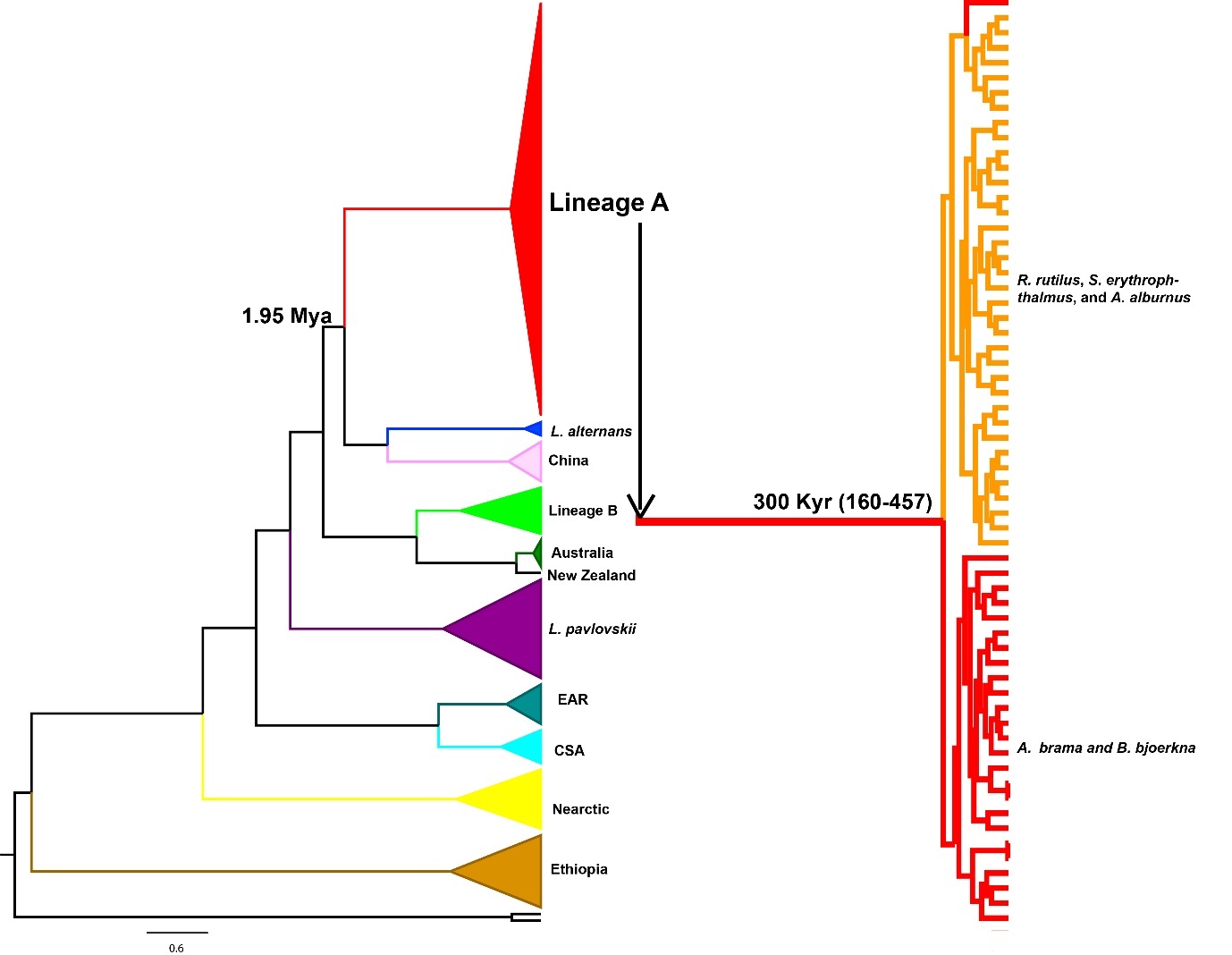

**Figure S1.** Dated phylogenetic tree derived from ddRAD sequence data, adapted from Nazarizadeh et al. 2023. The tree illustrates the evolutionary relationships among parasite populations. Lineage A, depicted in red, indicates two distinct sub-lineages: one found in *Abramis brama* and *Blicca bjoerkna*, and the other in *Rutilus rutilus*, *Scardinius erythrophthalmus*, and *Alburnus alburnus*. These sub-lineages diverged approximately 300 thousand years ago (Kyr), with the base of Lineage A tracing back to 1.95 million years ago (Mya)

**Table S1.** Detailed sampling information for the present study.

| Sample ID | Host | Locality | Country | Number of ddRAD reads | Barcode | Index | Accession number | Reference |
| --- | --- | --- | --- | --- | --- | --- | --- | --- |
| Li3_L1 | *Blicca bjoerkna* | Lipno reservoir | Czechia | 5486672 | CGATC | ATCACG | SAMN32032368 | Nazarizadeh et al., 2023 |
| Li5_L1 | *Blicca bjoerkna* | Lipno reservoir | Czechia | 5576052 | TCGAT | ATCACG | SAMN32032369 |  |
| Li9_L1 | *Abramis brama* | Lipno reservoir | Czechia | 5571409 | TGCAT | ATCACG | SAMN32032370 |  |
| Li12_L1 | *Abramis brama* | Lipno reservoir | Czechia | 4818508 | GGTTG | ATCACG | SAMN32032371 |  |
| S2_L1 | *Esox lucius* | Medard reservoir | Czechia | 6082925 | CGAAT | ATCACG | SAMN32032372 |  |
| CF_L1 | *Blicca bjoerkna* | Medard reservoir | Czechia | 6221424 | CGGCT | ATCACG | SAMN32032373 |  |
| PA1_L1 | *Rutilus rutilus* | Medard reservoir | Czechia | 4733833 | CGGTA | ATCACG | SAMN32032374 |  |
| PE1_L1 | *Rutilus rutilus* | Medard reservoir | Czechia | 6591323 | CGTAC | ATCACG | SAMN32032375 |  |
| CE2_L3 | *Blicca bjoerkna* | Medard reservoir | Czechia | 10733223 | TCACG | CGATGT | SAMN32032398 |  |
| PK_L3 | *Rutilus rutilus* | Medard reservoir | Czechia | 9863929 | TCCGG | CGATGT | SAMN32032399 |  |
| FR75_L4 | *Rutilus rutilus* | Créteil | France | 7353833 | GCATG | TTAGGC | SAMN32032400 |  |
| IE1_L4 | *Rutilus rutilus* | Lough Neagh | Ireland | 5816625 | TCGAT | TTAGGC | SAMN32032403 |  |
| IE2_L4 | *Rutilus rutilus* | Lough Neagh | Ireland | 5905188 | TGCAT | TTAGGC | SAMN32032404 |  |
| E2b_L4 | *Abramis brama* | Peipsi | Estonia | 5157082 | CATAT | TTAGGC | SAMN32032416 |  |
| TS04.126f_L4 | *Abramis brama* | Rybinsk | Russia | 4826446 | CGGCT | TTAGGC | SAMN32032418 |  |
| TS04.126h_L4 | *Abramis brama* | Rybinsk | Russia | 6194506 | CGTAC | TTAGGC | SAMN32032420 |  |
| TS04.126a_L4 | *Abramis brama* | Rybinsk | Russia | 4163635 | CGTCG | TTAGGC | SAMN32032421 |  |
| TS04.126c_L4 | *Abramis brama* | Rybinsk | Russia | 5453257 | CTGCG | TTAGGC | SAMN32032423 |  |
| UA1_L4 | *Alburnus alburnus* | Dniester | Ukraine | 6421321 | GTAGT | TTAGGC | SAMN32032433 |  |
| UA2_L4 | *Carassius carassius* | Dniester | Ukraine | 5673661 | GTCCG | TTAGGC | SAMN32032434 |  |
| UA3_L4 | *Rutilus rutilus* | Dniester | Ukraine | 7927417 | GTCGA | TTAGGC | SAMN32032435 |  |
| CR22_L4 | *Rutilus rutilus* | Créteil | France | 7761401 | TACCG | TTAGGC | SAMN32032436 |  |
| CR23_L4 | *Rutilus rutilus* | Créteil | France | 6568657 | TACGT | TTAGGC | SAMN32032437 |  |
| CR24_L4 | *Rutilus rutilus* | Créteil | France | 6807948 | TAGTA | TTAGGC | SAMN32032438 |  |
| CR25_L4 | *Rutilus rutilus* | Créteil | France | 6941629 | TATAC | TTAGGC | SAMN32032439 |  |
| CR26_L4 | *Rutilus rutilus* | Créteil | France | 6179618 | TCACG | TTAGGC | SAMN32032440 |  |
| X20M48 | *Rutilus rutilus* | Most | Czechia | 6035246 | CATAT | CGATGT | SAMN32032441 |  |
| LIPC1 | *Abramis brama* | Lipno | Czechia | 6191387 | CGGTA | CGATGT | SAMN32032444 |  |
| X20M57 | *Scardinius* | Most | Czechia | 6310454 | CTGAT | CGATGT | SAMN32032447 |  |
|  | *erythrophthalmus* |  |  |  |  |  |  |  |
| FR77 | *Rutilus rutilus* | Créteil | France | 4532929 | GCATG | ATCACG | SAMN32032463 |  |
| FR88 | *Rutilus rutilus* | Créteil | France | 6005983 | AACCA | ATCACG | SAMN32032464 |  |
| MPL28 | *Rutilus rutilus* | Most | Czechia | 6509615 | CGATC | ATCACG | SAMN32032465 |  |
| FR84 | *Rutilus rutilus* | Créteil | France | 5568505 | TCGAT | ATCACG | SAMN32032466 |  |
| FR89 | *Rutilus rutilus* | Créteil | France | 4560249 | TGCAT | ATCACG | SAMN32032467 |  |
| MPL29 | *Rutilus rutilus* | Most | Czechia | 5019649 | CAACC | ATCACG | SAMN32032468 |  |
| MPL30 | *Rutilus rutilus* | Most | Czechia | 4529673 | GGTTG | ATCACG | SAMN32032469 |  |
| X2Ab | *Abramis brama* | Římov | Czechia | 3418401 | AAGGA | ATCACG | SAMN32032470 |  |
| X3Ab | *Abramis brama* | Římov | Czechia | 4630225 | AGCTA | ATCACG | SAMN32032471 |  |
| X4Ab | *Abramis brama* | Římov | Czechia | 3987628 | ACACA | ATCACG | SAMN32032472 |  |
| X9Rr | *Rutilus rutilus* | Římov | Czechia | 4100447 | AATTA | ATCACG | SAMN32032473 |  |
| X16Ab | *Abramis brama* | Římov | Czechia | 3375762 | ACGGT | ATCACG | SAMN32032474 |  |
| Li2 | *Blicca bjoerkna* | Lipno | Czechia | 5036084 | CGGTA | ATCACG | SAMN32032483 |  |
| Li14 | *Abramis brama* | Lipno | Czechia | 5286573 | CGTAC | ATCACG | SAMN32032484 |  |
| X20R1PL | *Rutilus rutilus* | Římov | Czechia | 5850754 | CGTCG | ATCACG | SAMN32032485 |  |
| AB1 | *Blicca bjoerkna* | Římov | Czechia | 5028862 | CTGAT | ATCACG | SAMN32032486 |  |
| AB2 | *Blicca bjoerkna* | Římov | Czechia | 4445081 | CTGCG | ATCACG | SAMN32032487 |  |
| AbLipno1 | *Abramis brama* | Lipno | Czechia | 5418712 | GAGAT | ATCACG | SAMN32032488 |  |
| AbLipno2 | *Abramis brama* | Lipno | Czechia | 5094581 | GAGTC | ATCACG | SAMN32032489 |  |
| X20M11 | *Rutilus rutilus* | Most | Czechia | 5084222 | GCCGT | ATCACG | SAMN32032490 |  |
| X20M34 | *Rutilus rutilus* | Most | Czechia | 5876404 | GCTGA | ATCACG | SAMN32032491 |  |
| X20M51 | *Rutilus rutilus* | Most | Czechia | 4112140 | GTAGT | ATCACG | SAMN32032492 |  |
| X20M58 | *Scardinius* | Most | Czechia | 5502583 | GTCCG | ATCACG | SAMN32032493 |  |
|  | *erythrophthalmus* |  |  |  |  |  |  |  |
| Ea2 | *Abramis brama* | Peipsi | Estonia | 4480806 | TAGTA | ATCACG | SAMN32032496 |  |
| MPR1 | *Scardinius* | Most | Czechia | 6283789 | TCAGT | ATCACG | SAMN32032498 |  |
|  | *erythrophthalmus* |  |  |  |  |  |  |  |
| MPR2 | *Scardinius* | Most | Czechia | 5779666 | TCCGG | ATCACG | SAMN32032499 |  |
|  | *erythrophthalmus* |  |  |  |  |  |  |  |
| MPR3 | *Scardinius* | Most | Czechia | 5855323 | TCTGC | ATCACG | SAMN32032500 |  |
|  | *erythrophthalmus* |  |  |  |  |  |  |  |
| MPL27B | *Rutilus rutilus* | Most | Czechia | 5807870 | TTACC | ATCACG | SAMN32032502 |  |
| Ir8_L3 | *Squalius orientalis* |  | Iran | 8054152 | GCTGA | CGATGT | SAMN32032506 |  |
| A14.1_L2 | *Abramis brama* | Žermanice | Czechia | 4468390 | CGTCG | ATCACG | xxxxxx | Present Study |
| A14.2_L2 | *Abramis brama* | Žermanice | Czechia | 4489443 | CTGAT | ATCACG | xxxxxx |  |
| A3.1_L2 | *Abramis brama* | Těrlicko | Czechia | 12020491 | GAGTC | ATCACG | xxxxxx |  |
| A3.2_L2 | *Abramis brama* | Těrlicko | Czechia | 13884205 | GCCGT | ATCACG | xxxxxx |  |
| A3.3_L3 | *Abramis brama* | Těrlicko | Czechia | 13128679 | GCTGA | ATCACG | xxxxxx |  |
| A3.4_L3 | *Abramis brama* | Těrlicko | Czechia | 11289209 | GGATA | ATCACG | xxxxxx |  |
| A3.6_L3 | *Abramis brama* | Těrlicko | Czechia | 16639212 | GGCCA | ATCACG | xxxxxx |  |
| A3.7_L3 | *Abramis brama* | Těrlicko | Czechia | 14310464 | GGCTC | ATCACG | xxxxxx |  |
| A8.2_L3 | *Abramis brama* | Těrlicko | Czechia | 14870785 | GTAGT | ATCACG | xxxxxx |  |
| Li12_L1 | *Abramis brama* | Lipno | Czechia | 14473172 | TCAGT | ATCACG | xxxxxx |  |
| Li14 | *Abramis brama* | Lipno | Czechia | 14527049 | TCCGG | ATCACG | xxxxxx |  |
| Li15_L1 | *Abramis brama* | Lipno | Czechia | 15586050 | TCTGC | ATCACG | xxxxxx |  |
| Li9_L1 | *Abramis brama* | Lipno | Czechia | 8774995 | ACTTC | CGATGT | xxxxxx |  |
| LIPC1 | *Abramis brama* | Lipno | Czechia | 6889676 | ATACG | CGATGT | xxxxxx |  |
| LIPC1A | *Abramis brama* | Lipno | Czechia | 8040507 | ATGAG | CGATGT | xxxxxx |  |
| A11.1_L3 | *Alburnus alburnus* | Žermanice | Czechia | 5094581 | GAGTC | ATCACG | xxxxxx |  |
| A13.1_L2 | *Alburnus alburnus* | Žermanice | Czechia | 5084222 | GCCGT | ATCACG | xxxxxx |  |
| A2.2_L3 | *Alburnus alburnus* | Těrlicko | Czechia | 5876404 | GCTGA | ATCACG | xxxxxx |  |
| A2.3_L2 | *Alburnus alburnus* | Těrlicko | Czechia | 4112140 | GTAGT | ATCACG | xxxxxx |  |
| A2.5_L3 | *Alburnus alburnus* | Těrlicko | Czechia | 5502583 | GTCCG | ATCACG | xxxxxx |  |
| A2.5_L4 | *Alburnus alburnus* | Těrlicko | Czechia | 4480806 | TAGTA | ATCACG | xxxxxx |  |
| A2.8_L2 | *Alburnus alburnus* | Těrlicko | Czechia | 6283789 | TCAGT | ATCACG | xxxxxx |  |
| A20.1_L2 | *Alburnus alburnus* | Žermanice | Czechia | 5779666 | TCCGG | ATCACG | xxxxxx |  |
| A20.2_L2 | *Alburnus alburnus* | Žermanice | Czechia | 5855323 | TCTGC | ATCACG | xxxxxx |  |
| A23.1_L3 | *Alburnus alburnus* | Žermanice | Czechia | 5807870 | TTACC | ATCACG | xxxxxx |  |
| A23.2_L3 | *Alburnus alburnus* | Žermanice | Czechia | 8054152 | GCTGA | CGATGT | xxxxxx |  |
| A9.1_L2 | *Alburnus alburnus* | Žermanice | Czechia | 4468390 | CGTCG | ATCACG | xxxxxx |  |
| CZ77 | *Alburnus alburnus* | Želivka | Czechia | 4489443 | CTGAT | ATCACG | xxxxxx |  |
| CZ81 | *Alburnus alburnus* | Želivka | Czechia | 12020491 | GAGTC | ATCACG | xxxxxx |  |
| CZ82 | *Alburnus alburnus* | Želivka | Czechia | 13884205 | GCCGT | ATCACG | xxxxxx |  |
| CZ83 | *Alburnus alburnus* | Želivka | Czechia | 13128679 | GCTGA | ATCACG | xxxxxx |  |
| CZ84 | *Alburnus alburnus* | Želivka | Czechia | 11289209 | GGATA | ATCACG | xxxxxx |  |
| CZ86 | *Alburnus alburnus* | Želivka | Czechia | 16639212 | GGCCA | ATCACG | xxxxxx |  |
| X20LIPO19 | *Alburnus alburnus* | Lipno | Czechia | 14310464 | GGCTC | ATCACG | xxxxxx |  |
| X20LIPO3 | *Alburnus alburnus* | Lipno | Czechia | 14870785 | GTAGT | ATCACG | xxxxxx |  |
| CNA_L1 | *Blicca bjoerkna* | Medard | Czechia | 14497443 | TACGT | ATCACG | xxxxxx |  |
| CND_L1 | *Blicca bjoerkna* | Medard | Czechia | 15006962 | TAGTA | ATCACG | xxxxxx |  |
| JO1_L1 | *Blicca bjoerkna* | Jordan | Czechia | 14473172 | TCAGT | ATCACG | xxxxxx |  |
| Li1_L1 | *Blicca bjoerkna* | Lipno | Czechia | 14527049 | TCCGG | ATCACG | xxxxxx |  |
| Li1_L3 | *Blicca bjoerkna* | Lipno | Czechia | 15586050 | TCTGC | ATCACG | xxxxxx |  |
| A1.1_L2 | *Rutilus rutilus* | Těrlicko | Czechia | 8081375 | ATTAC | CGATGT | xxxxxx |  |
| A1.2_L2 | *Rutilus rutilus* | Těrlicko | Czechia | 6861512 | CATAT | CGATGT | xxxxxx |  |
| A1.3_L3 | *Rutilus rutilus* | Těrlicko | Czechia | 8609554 | CGGTA | CGATGT | xxxxxx |  |
| A1.4_L2 | *Rutilus rutilus* | Těrlicko | Czechia | 5418712 | GAGAT | ATCACG | xxxxxx |  |
| A22.1_L3 | *Rutilus rutilus* | Žermanice | Czechia | 5094581 | GAGTC | ATCACG | xxxxxx |  |
| A6.1_L3 | *Rutilus rutilus* | Těrlicko | Czechia | 5084222 | GCCGT | ATCACG | xxxxxx |  |
| A6.2_L3 | *Rutilus rutilus* | Těrlicko | Czechia | 5876404 | GCTGA | ATCACG | xxxxxx |  |
| A6.3_L3 | *Rutilus rutilus* | Těrlicko | Czechia | 4112140 | GTAGT | ATCACG | xxxxxx |  |
| A6.4_L3 | *Rutilus rutilus* | Těrlicko | Czechia | 5502583 | GTCCG | ATCACG | xxxxxx |  |
| A7.1_L3 | *Rutilus rutilus* | Těrlicko | Czechia | 4480806 | TAGTA | ATCACG | xxxxxx |  |
| CZ90 | *Rutilus rutilus* | Želivka | Czechia | 6283789 | TCAGT | ATCACG | xxxxxx |  |
| CZ92 | *Rutilus rutilus* | Želivka | Czechia | 5779666 | TCCGG | ATCACG | xxxxxx |  |
| CZ94 | *Rutilus rutilus* | Želivka | Czechia | 5855323 | TCTGC | ATCACG | xxxxxx |  |
| CZ95 | *Rutilus rutilus* | Želivka | Czechia | 5807870 | TTACC | ATCACG | xxxxxx |  |
| CZ96 | *Rutilus rutilus* | Želivka | Czechia | 8054152 | GCTGA | CGATGT | xxxxxx |  |
| KL1_L1 | *Rutilus rutilus* | Klíčava | Czechia | 4468390 | CGTCG | ATCACG | xxxxxx |  |
| KL1_L3 | *Rutilus rutilus* | Klíčava | Czechia | 4489443 | CTGAT | ATCACG | xxxxxx |  |
| Li11B_L1 | *Rutilus rutilus* | Lipno | Czechia | 12020491 | GAGTC | ATCACG | xxxxxx |  |
| N1 | *Rutilus rutilus* | Nýrsko | Czechia | 14310464 | GGCTC | ATCACG | xxxxxx |  |
| N2 | *Rutilus rutilus* | Nýrsko | Czechia | 14870785 | GTAGT | ATCACG | xxxxxx |  |
| N3 | *Rutilus rutilus* | Nýrsko | Czechia | 13449952 | GTCCG | ATCACG | xxxxxx |  |
| N4 | *Rutilus rutilus* | Nýrsko | Czechia | 16996354 | GTCGA | ATCACG | xxxxxx |  |
| N8 | *Rutilus rutilus* | Nýrsko | Czechia | 14497443 | TACGT | ATCACG | xxxxxx |  |
| NPL1_L3 | *Rutilus rutilus* | Nýrsko | Czechia | 15006962 | TAGTA | ATCACG | xxxxxx |  |
| NPL10A_L3 | *Rutilus rutilus* | Nýrsko | Czechia | 14473172 | TCAGT | ATCACG | xxxxxx |  |
| NPL2_L3 | *Rutilus rutilus* | Nýrsko | Czechia | 14527049 | TCCGG | ATCACG | xxxxxx |  |
| NPL3_L3 | *Rutilus rutilus* | Nýrsko | Czechia | 15586050 | TCTGC | ATCACG | xxxxxx |  |
| NPL4_L3 | *Rutilus rutilus* | Nýrsko | Czechia | 8774995 | ACTTC | CGATGT | xxxxxx |  |
| NPL6_L3 | *Rutilus rutilus* | Nýrsko | Czechia | 6889676 | ATACG | CGATGT | xxxxxx |  |
| NPL7_L3 | *Rutilus rutilus* | Nýrsko | Czechia | 8040507 | ATGAG | CGATGT | xxxxxx |  |
| NPL9_L3 | *Rutilus rutilus* | Nýrsko | Czechia | 8081375 | ATTAC | CGATGT | xxxxxx |  |
| NPL9_L4 | *Rutilus rutilus* | Nýrsko | Czechia | 6861512 | CATAT | CGATGT | xxxxxx |  |
| P10 | *Rutilus rutilus* | Medard | Czechia | 8609554 | CGGTA | CGATGT | xxxxxx |  |
| P2 | *Rutilus rutilus* | Medard | Czechia | 5418712 | GAGAT | ATCACG | xxxxxx |  |
| PB_L2 | *Rutilus rutilus* | Medard | Czechia | 5084222 | GCCGT | ATCACG | xxxxxx |  |
| PE_L1 | *Rutilus rutilus* | Medard | Czechia | 5876404 | GCTGA | ATCACG | xxxxxx |  |
| PH_L2 | *Rutilus rutilus* | Medard | Czechia | 5502583 | GTCCG | ATCACG | xxxxxx |  |
| PK_L2 | *Rutilus rutilus* | Medard | Czechia | 4480806 | TAGTA | ATCACG | xxxxxx |  |
| PQ_L2 | *Rutilus rutilus* | Medard | Czechia | 5779666 | TCCGG | ATCACG | xxxxxx |  |
| PX_L1 | *Rutilus rutilus* | Medard | Czechia | 5855323 | TCTGC | ATCACG | xxxxxx |  |
| X20M40 | *Rutilus rutilus* | Most | Czechia | 4468390 | CGTCG | ATCACG | xxxxxx |  |
| X20M44 | *Rutilus rutilus* | Most | Czechia | 4489443 | CTGAT | ATCACG | xxxxxx |  |
| X20M48A | *Rutilus rutilus* | Most | Czechia | 13884205 | GCCGT | ATCACG | xxxxxx |  |
| ZE1_L1 | *Rutilus rutilus* | Želivka | Czechia | 14310464 | GGCTC | ATCACG | xxxxxx |  |
| ZE2_L1 | *Rutilus rutilus* | Želivka | Czechia | 14870785 | GTAGT | ATCACG | xxxxxx |  |
| ZE2_L3 | *Rutilus rutilus* | Želivka | Czechia | 13449952 | GTCCG | ATCACG | xxxxxx |  |
| A5.1_L3 | *Rutilus rutilus* | Těrlicko | Czechia | 15586050 | TCTGC | ATCACG | xxxxxx |  |

**Table S2.** Comparison of different divergence scenarios, analysed using fastsimcoal2, for two parasite populations associated with RSA and AB hosts.

| **Model Parameters** | **Search range (log uniform distribution)** | **Allopatry** | **primary contact** | **Secondary contact** | **Isolation with continuous gene flow** |
| --- | --- | --- | --- | --- | --- |
| **RSA** | 100-100000 | 11,561 | 7,070 | 7,861 | **5,211** |
| **AB** | 100-100000 | 48,323 | 5,541 | 4,935 | **5,332** |
| **MRCA (LineageA)** | resize 0.1-10 | 11,549 | 3,723 | 3,029 | **3,627** |
| **Time** | 1- 300,000 | NA | 60000 | 80000 | **120,000** |
| **early mig RSA->AB** | 1x10^-6^-0.01 | NA | NA | 3.3x10^-4^/1.64 | **4.2x10^-6^/0.02** |
| **early mig AB->RSA** | 1x10^-6^-0.01 | NA | NA | 6.8x10^-4^/4.00 | **3.3x10^-4^/1.96** |
| **recent mig RSA->AB** | 1x10^-6^-0.01 | NA | 4.4x10^-4^/2.42 | NA | **9.4x10^-4^/5.00** |
| **recent mig AB->RSA** | 1x10^-6^-0.01 | NA | 9.4x10^-4^/6.66 | NA | **7.1x10^-4^/4.22** |
| Log10 |  | -2144.5 | -1706.8 | -1783.9 | **-1623.1** |
| DAIC |  | 96.4 | 25.4 | 28.1 | **0.00** |
| AIC |  | 4367.2 | 3821.6 | 3867.5 | **3150.2** |
| AIC_w |  | 0.00 | 0.19 | 0.14 | **0.81** |

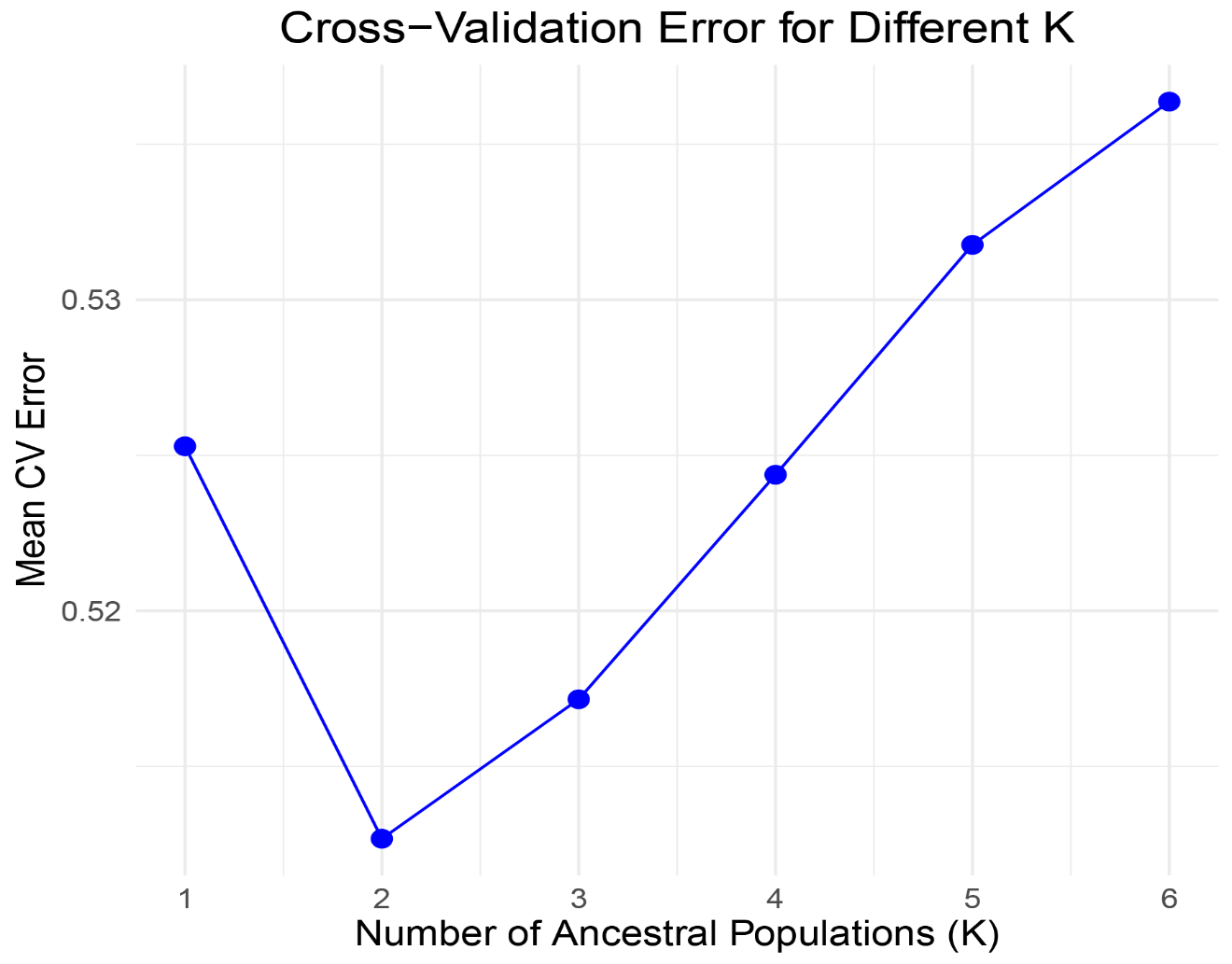

**Figure S2.** Cross-validation error rates from admixture analysis indicating the presence of two distinct genetic clusters among the parasite populations studied. The analysis assesses the likelihood of various numbers of genetic clusters (K), with the lowest cross-validation error suggesting the most probable number of clusters. In this case, the figure highlights that a model with two clusters (K=2) best fits the data, pointing to a significant genetic structuring within the parasite populations.

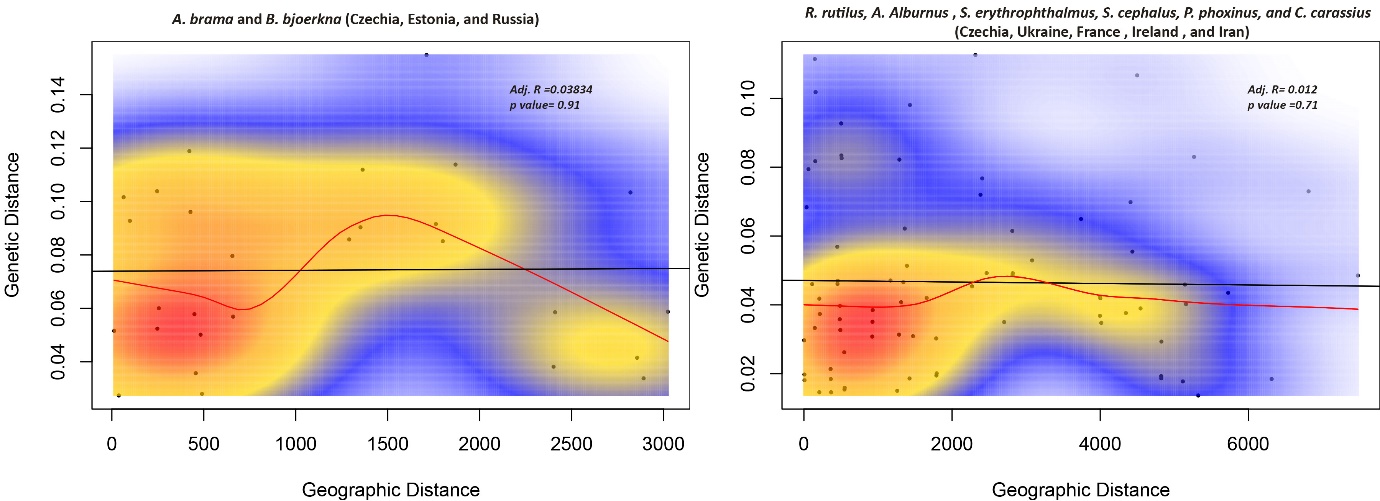

**Figure S3.** Comparative Mantel tests illustrating the lack of significant correlation between genetic distance and geographic distance in two parasite genetic clusters: one within Abramis brama and Blicca bjoerkna populations across Czechia, Estonia, and Russia (left panel), and another within *Rutilus rutilus, Scardinius erythrophthalmus, Alburnus alburnus, Phoxinus phoxinus*, and *Carassius carassius* populations across Czechia, Ukraine, France, Ireland, and Iran (right panel). Both analyses reveal low and nonsignificant adjusted R-values, indicating that geographic separation does not predict genetic differentiation within these clusters.

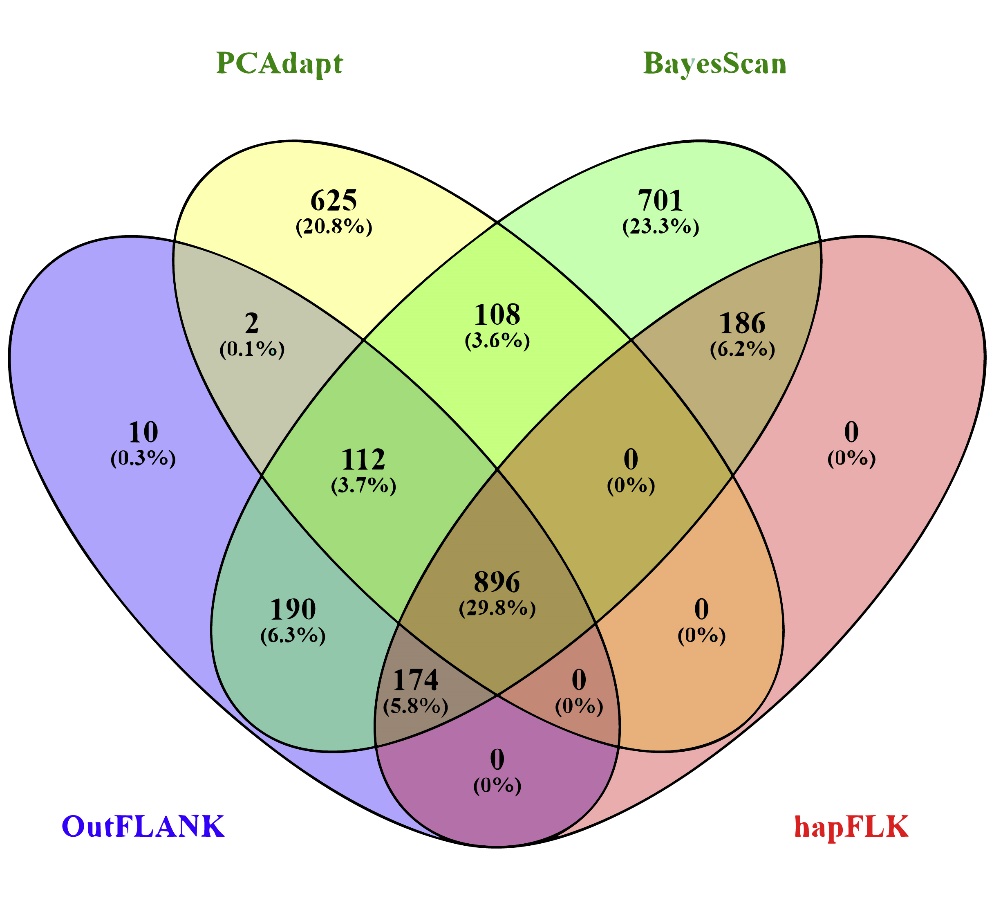

**Figure S4.** Venn diagram depicting the overlap of genome-wide SNP analyses using four different selection analysis methods: PCAadapt, BayeScan, OutFLANK, and hapFLK. The intersection at the center represents 896 SNPs (29.8% of the total) identified as under selection by all methods, highlighting a consensus in the detection of selective pressures across the genome.

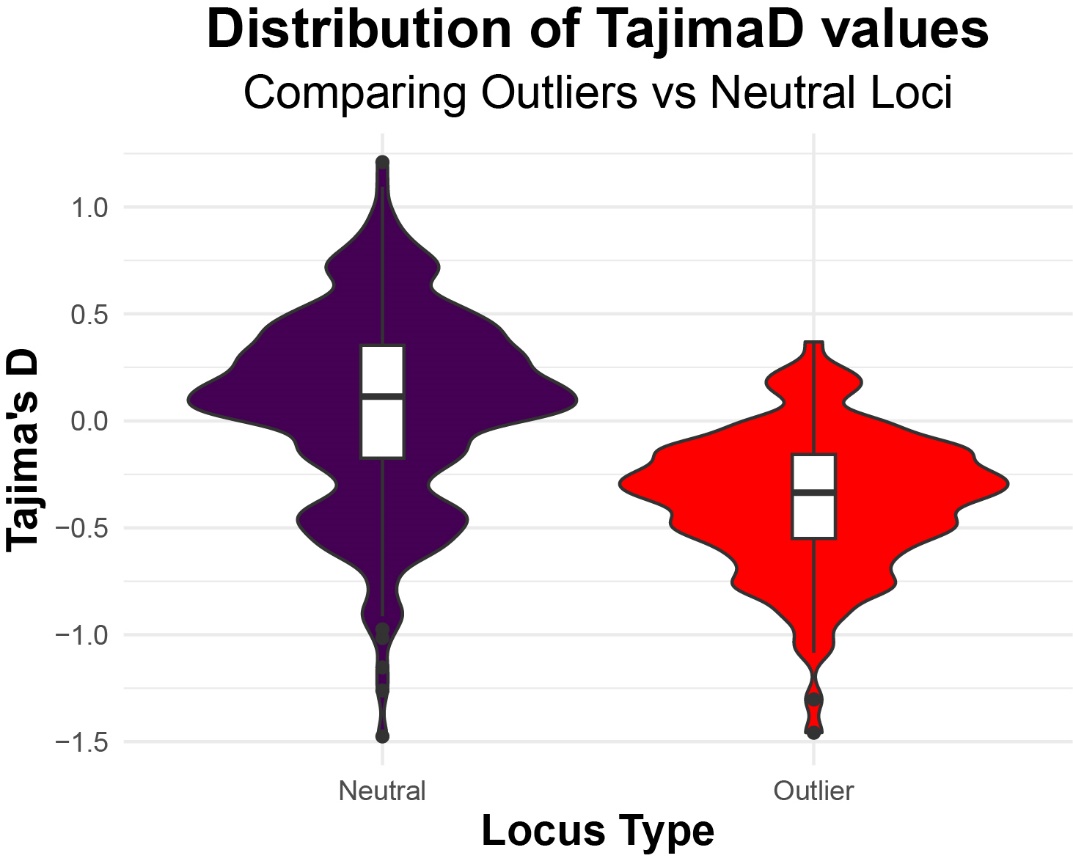

**Figure S5** Violin plot contrasting the distribution of Tajima's D values between neutral loci (purple) and outlier loci (red). The data show that outlier loci have a significantly more negative distribution of Tajima's D values, suggesting a deviation from neutrality, potentially due to selective sweeps.

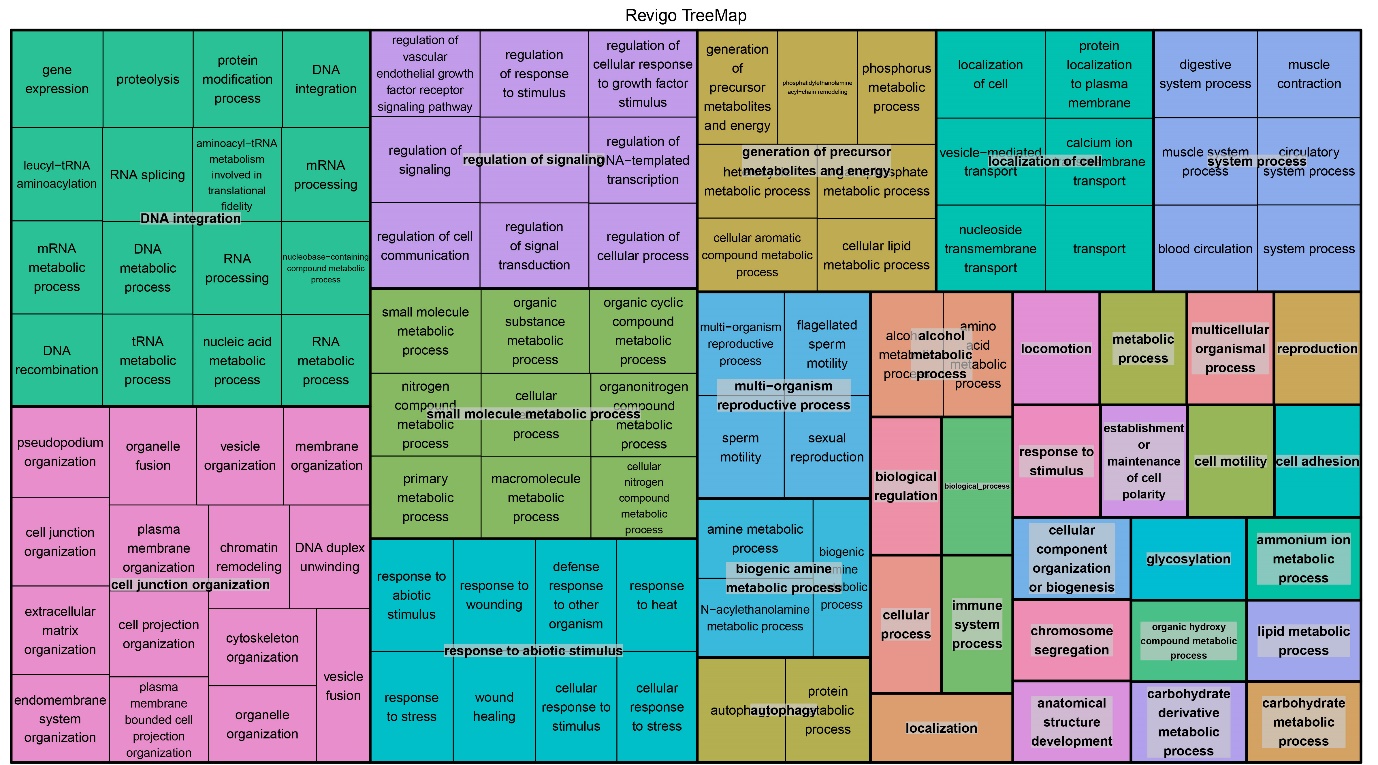

**Figure S6.** Revigo TreeMap visualization of the functional annotation of SNPs under selection, categorized by biological processes. Each colored block represents a unique cluster of related biological functions, with size indicating the frequency of the process in the dataset. This map highlights the diverse biological processes potentially influenced by selective pressures.

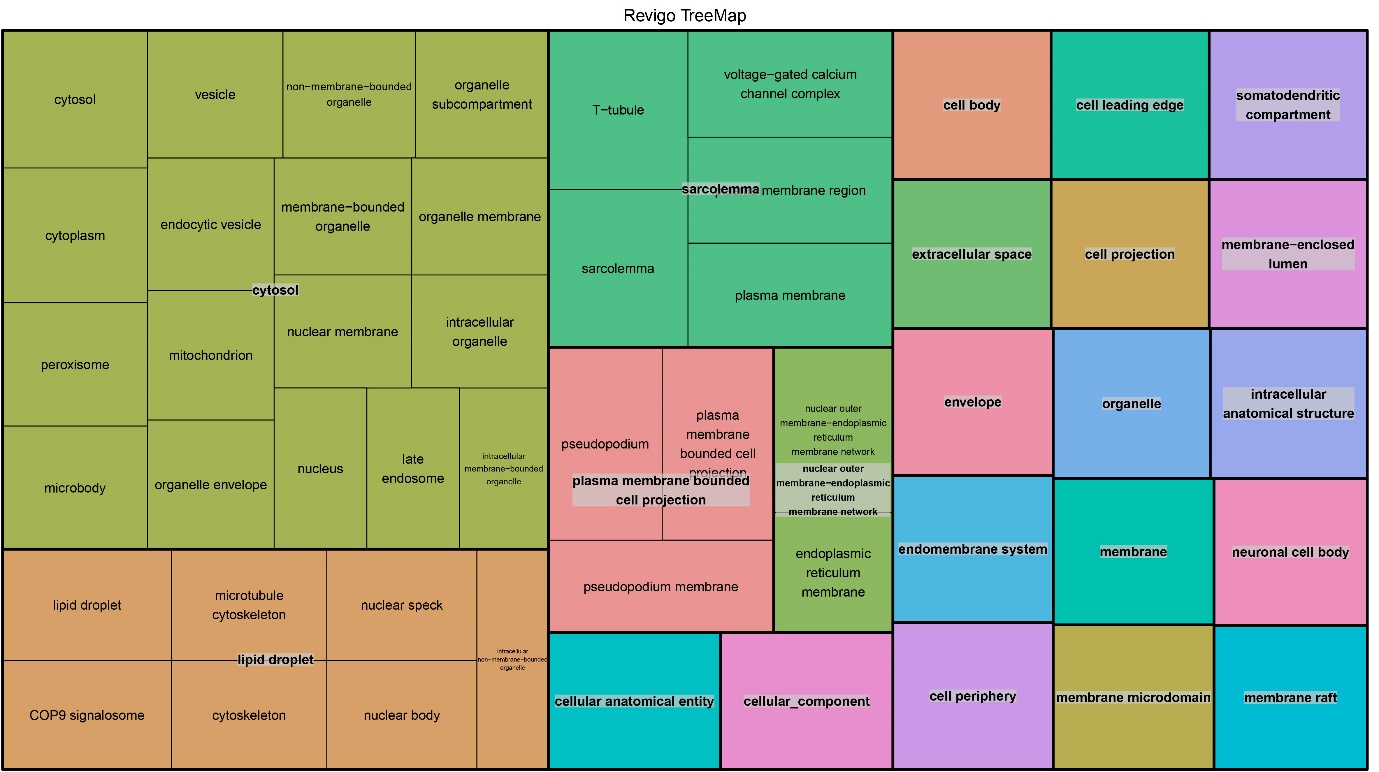

Figure S7 Revigo TreeMap visualization representing the cellular component categorization of SNPs under selection. Each block signifies a distinct cellular component, with size corresponding to the prevalence of SNPs associated with that component in the data. This distribution showcases the complexity of cellular architecture impacted by selection, ranging from broader components like the cytosol and plasma membrane to specific structures such as lipid droplets and organelle membranes

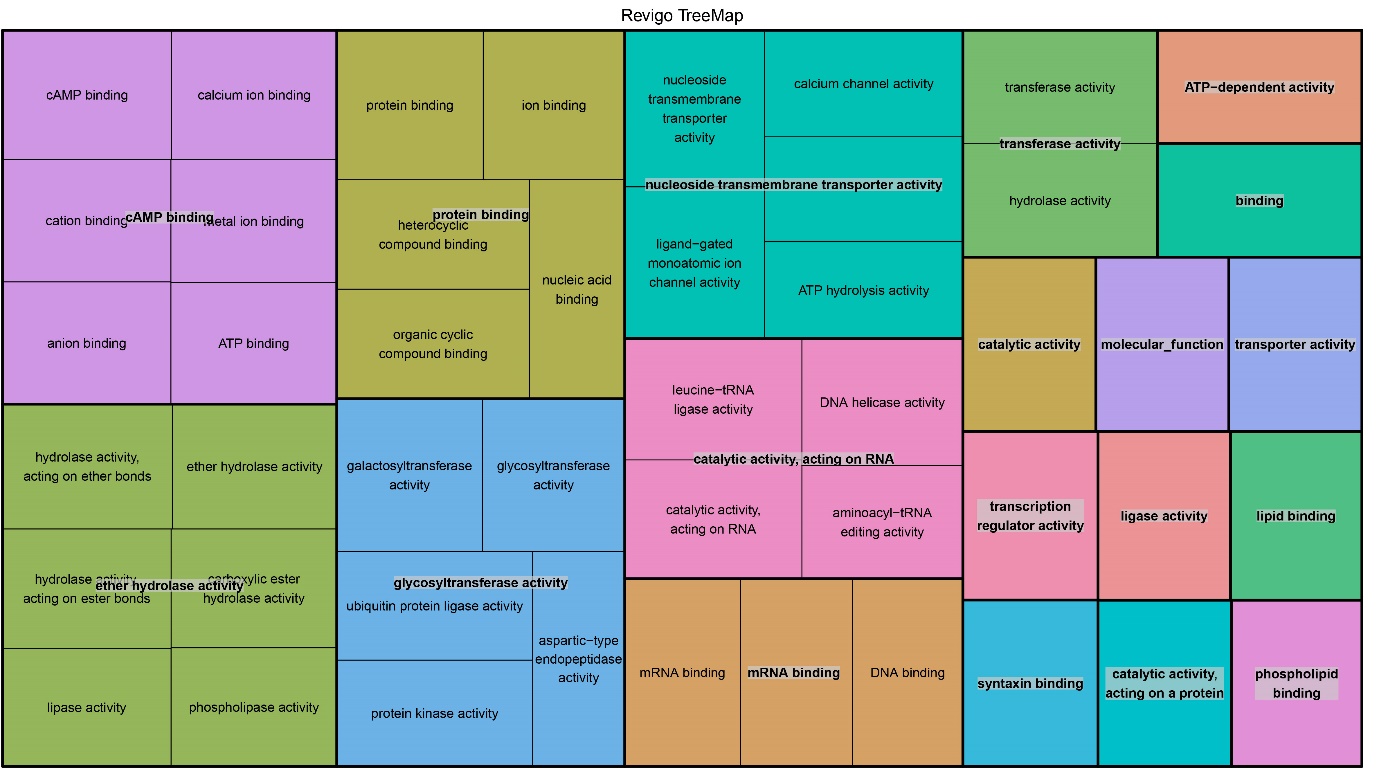

**Figure S8.** Revigo TreeMap visualization mapping the molecular function categorization of SNPs under selection. Each block represents a distinct molecular function, with size proportional to the occurrence of SNPs associated with that function in the dataset. The map illustrates the intricate network of molecular activities, from binding and catalytic functions to transport and enzyme activities, influenced by selective pressures at the molecular level.
